# Rescue of behavioral and electrophysiological phenotypes in a Pitt-Hopkins syndrome mouse model by genetic restoration of Tcf4 expression

**DOI:** 10.1101/2021.08.02.454783

**Authors:** Hyojin Kim, Eric B. Gao, Adam Draper, Noah C. Berens, Hanna Vihma, Xinyuan Zhang, Alexandra Higashi-Howard, Kimberly D. Ritola, Jeremy M. Simon, Andrew J. Kennedy, Benjamin D. Philpot

## Abstract

Pitt-Hopkins syndrome (PTHS) is a neurodevelopmental disorder caused by monoallelic mutation or deletion in the *transcription factor 4* (*TCF4*) gene. Individuals with PTHS typically present in the first year of life with developmental delay and exhibit intellectual disability, lack of speech, and motor incoordination. There are no effective treatments available for PTHS, but the root cause of the disorder, *TCF4* haploinsufficiency, suggests that it could be treated by normalizing *TCF4* gene expression. Here we performed proof-of-concept viral gene therapy experiments using a conditional *Tcf4* mouse model of PTHS and found that postnatally reinstating *Tcf4* expression in neurons improved anxiety-like behavior, activity levels, innate behaviors, and memory. Postnatal reinstatement also partially corrected EEG abnormalities, which we characterized here for the first time, and the expression of key TCF4-regulated genes. Our results support a genetic normalization approach as a treatment strategy for PTHS, and possibly other TCF4-linked disorders.

## Introduction

Pitt-Hopkins syndrome (PTHS) is a severe neurodevelopmental disorder, characterized by delay in motor function, lack of speech, stereotypies, sleep disorder, seizures, and intellectual disability. Other commonly reported features include constipation and hyperventilation (Bedeschi et al., 2017; Goodspeed et al., 2018; Zollino et al., 2019). While PTHS is a lifelong disorder, there are currently no treatments for PTHS (Zollino et al., 2019). PTHS is caused by monoallelic mutation or deletion of *transcription factor 4* (*TCF4*), which is a member of the class I basic-helix-loop-helix (bHLH) group (Amiel et al., 2007; Zweier et al., 2007). PTHS- causing mutations typically impair the function of the bHLH domain (Giurgea et al., 2008; Sepp, Pruunsild, & Timmusk, 2012), which is responsible for dimerizing with other bHLH proteins and for binding to Ephrussi box DNA elements to regulate transcription (Dennis, Han, & Schuurmans, 2019). Thus, targeting genes dysregulated by *TCF4* haploinsufficiency could potentially serve as a therapeutic intervention. However, hundreds to thousands of genes lie downstream of TCF4 (Doostparast Torshizi et al., 2019; M. P. Forrest, Waite, Martin-Rendon, & Blake, 2013; Hill et al., 2017; Xia et al., 2018), making it nearly impossible to find transcriptional modifiers to correct their expression levels. While targeting TCF4-impacted genes presents a fundamental conceptual challenge as a therapeutic approach, directly overcoming the core genetic defect underlying PTHS may offer a more effective treatment strategy.

In principle, PTHS phenotypes could be prevented or corrected by normalizing *TCF4* expression, with the degree of efficacy likely impacted by the age and specificity of the intervention. Convergent lines of evidence support the idea that the disorder can be treated, at least to a degree, throughout life. For example, studies in animal models of other single-gene neurodevelopmental disorders, including Rett and Angelman syndromes, have shown that normalizing expression of the disease-causing gene in postnatal life could provide therapeutic benefits (Guy, Gan, Selfridge, Cobb, & Bird, 2007; Silva-Santos et al., 2015). Therefore, the same might be true for PTHS. While synaptic defects have been observed in mouse models of PTHS, there is no evidence for disease-related neurodegeneration in PTHS individuals or mouse models (Rannals et al., 2016; Thaxton et al., 2018). Therefore, the observed synaptic defects could be reversible. In support of this idea, subtle upregulation of *Tcf4* expression by knocking down *Hdac2* has been shown to partially rescue learning and memory in adult PTHS model mice (Kennedy et al., 2016). Collectively, these observations indicate that PTHS might benefit from genetic normalization approaches to compensate for loss-of-function of TCF4 such as gene therapy, antisense oligonucleotides (ASOs), and small molecules.

A critical question that must be addressed prior to developing genetic normalization approaches for PTHS is whether behavioral phenotypes can be rescued if *TCF4* expression is restored during postnatal development. This question is particularly intriguing given observations that *TCF4/Tcf4* expression in the human/mouse brain peaks perinatally, before subsequently declining to basal levels that are sustained throughout adulthood (Phan et al., 2020; Rannals et al., 2016). Here, we leveraged a conditional mouse model to establish the extent to which conditional reinstatement of *Tcf4* expression could rescue behavioral phenotypes in a mouse model of PTHS. We first validated our approach by demonstrating that embryonic pan-cellular reinstatement of *Tcf4* expression could fully prevent PTHS-associated phenotypes, while embryonic *Tcf4* reinstatement selectively in excitatory or inhibitory neurons led to rescue only a subset of behavioral phenotypes. We then modeled viral gene therapy to show that postnatal reinstatement of *Tcf4* expression in neurons can fully or partially rescue behavioral and electrophysiological phenotypes in a mouse model of PTHS. Our results provide evidence that postnatal genetic normalization strategies offer an effective therapeutic intervention for PTHS.

## Results

### Pan-cellular embryonic reinstatement of *Tcf4* fully rescues behavioral phenotypes in a PTHS mouse model

We generated a conditional *Tcf4* reinstatement mouse model of PTHS (*Tcf4^STOP/+^*) in which a STOP cassette and GFP reporter, flanked by loxP sites, were inserted upstream of the basic Helix-Loop-Helix (bHLH) DNA binding domain in exon 18 of *Tcf4* (**Figure 1A**). To produce embryonic, pan-cellular reinstatement of *Tcf4*, we crossed *Tcf4^STOP/+^* mice to transgenic mice expressing Cre under the β-Actin promoter (Jagle, Gasser, Muller, & Kinzel, 2007). As predicted from our design, the levels of full-length TCF4/*Tcf4* were reduced by half in *Tcf4^STOP/+^* mouse brain compared to *Tcf4^+/+^* (wildtype control) mouse brain and fully normalized by crossing *Tcf4^STOP/+^* to *Actin-Cre^+/-^* mice (**Figure 1B**, *Tcf4^+/+^* : 1.0 ± 0.07, n = 7; *Tcf4^STOP/+^* : 0.58 ± 0.05, n = 7; *Tcf4^STOP/+^::Actin-Cre* : 1.07 ± 0.07, n = 5, and **Figure 1-figure supplement 1A**, *Tcf4^+/+^* : 1.0 ± 0.09, n = 7; *Tcf4^STOP/+^* : 0.61 ± 0.07, n = 7; *Tcf4^STOP/+^::Actin-Cre* : 1.04 ± 0.09, n = 5). We stained for the GFP reporter in sagittal brain sections from *Tcf4^+/+^*, *Tcf4^STOP/+^*, and *Tcf4^STOP/+^::Actin-Cre* mice (**Figure 1-figure supplement 1B**). GFP signal, a proxy for presence of the STOP cassette, was detected throughout the *Tcf4^STOP/+^* mouse brain. By contrast, this signal was absent from *Tcf4^+/+^* and *Tcf4^STOP/+^::Actin-Cre* brain sections, indicating efficient excision of the GFP-STOP cassette and concomitant reinstatement of biallelic *Tcf4* expression in *Tcf4^STOP/+^::Actin-Cre* model (**Figure 1-figure supplement 1B**). To demonstrate the consequences of pan-cellular embryonic *Tcf4* reinstatement in PTHS model mice, we studied a variety of physiological functions and behaviors in control (*Tcf4^+/+^* and *Tcf4^+/+^::Actin-Cre*), PTHS model (*Tcf4^STOP/+^*), and reinstatement model (*Tcf4^STOP/+^::Actin-Cre*) mice. Male and female *Tcf4^STOP/+^* mice had reduced body and brain weights, whereas *Tcf4^STOP/+^::Actin-Cre* body and brain weights were similar to their littermate controls (**Figure 1-figure supplement 1C-D**). This suggests that embryonic reinstatement of *Tcf4* expression could prevent microcephaly in PTHS model mice. To study whether long-term memory deficits could be prevented, we examined object location memory by measuring interaction time of identical objects, with one object located in the familiar position and the other in a novel position. *Tcf4^STOP/+^* interactions with objects located in the familiar and novel positions were of similar duration, suggesting the inability to remember the previously-presented location of the object and suggestive of long-term memory deficits (**Figure 1C**). *Tcf4^STOP/+^::Actin-Cre* and control mice both interacted to a significantly greater extent with the object located in the novel position, suggesting recovery of normal long-term memory function subsequent to embryonic, pan-cellular *Tcf4* reinstatement (**Figure 1C**). We then assessed locomotor and exploration activity by the open field test and found that *Tcf4^STOP/+^* mice showed increased activity and total distance travelled compared to *Tcf4^STOP/+^::Actin-Cre* mice, which exhibited control levels of activity (**Figure 1D**). We also examined nest building, an innate, goal-directed behavior achieved by pulling, carrying, fraying, push digging, sorting, and fluffing of nest material (Deacon, 2006). *Tcf4^STOP/+^* mice exhibited poor performance in the nest building task over the 4-day testing period, using roughly half the nest material used by control mice. This phenotype was completely rescued in the *Tcf4^STOP/+^::Actin-Cre* model (**Figure 1E**). To assess anxiety-like behavior, we evaluated mice in the elevated plus maze task. We observed *Tcf4^STOP/+^* mice to spend similar amounts of time in the closed and open arms, indicating an apparent low-anxiety phenotype. This behavioral feature also appeared to be normalized in *Tcf4^STOP/+^::Actin-Cre* mice, spending proportionally more time in the closed arms (**Figure 1F**). Collectively, our results confirm that *Tcf4^STOP/+^* mice exhibit physiological and behavioral phenotypes like those observed in other mouse models of PTHS (Kennedy et al., 2016; Thaxton et al., 2018), demonstrating the efficacy of the transcriptional STOP cassette in blocking TCF4 function. Moreover, these data show that *Tcf4* reinstatement upon embryonic Cre-mediated excision of the STOP cassette can fully prevent the emergence of physiological and behavioral deficits associated with *Tcf4* haploinsufficiency.

**Figure 1.**
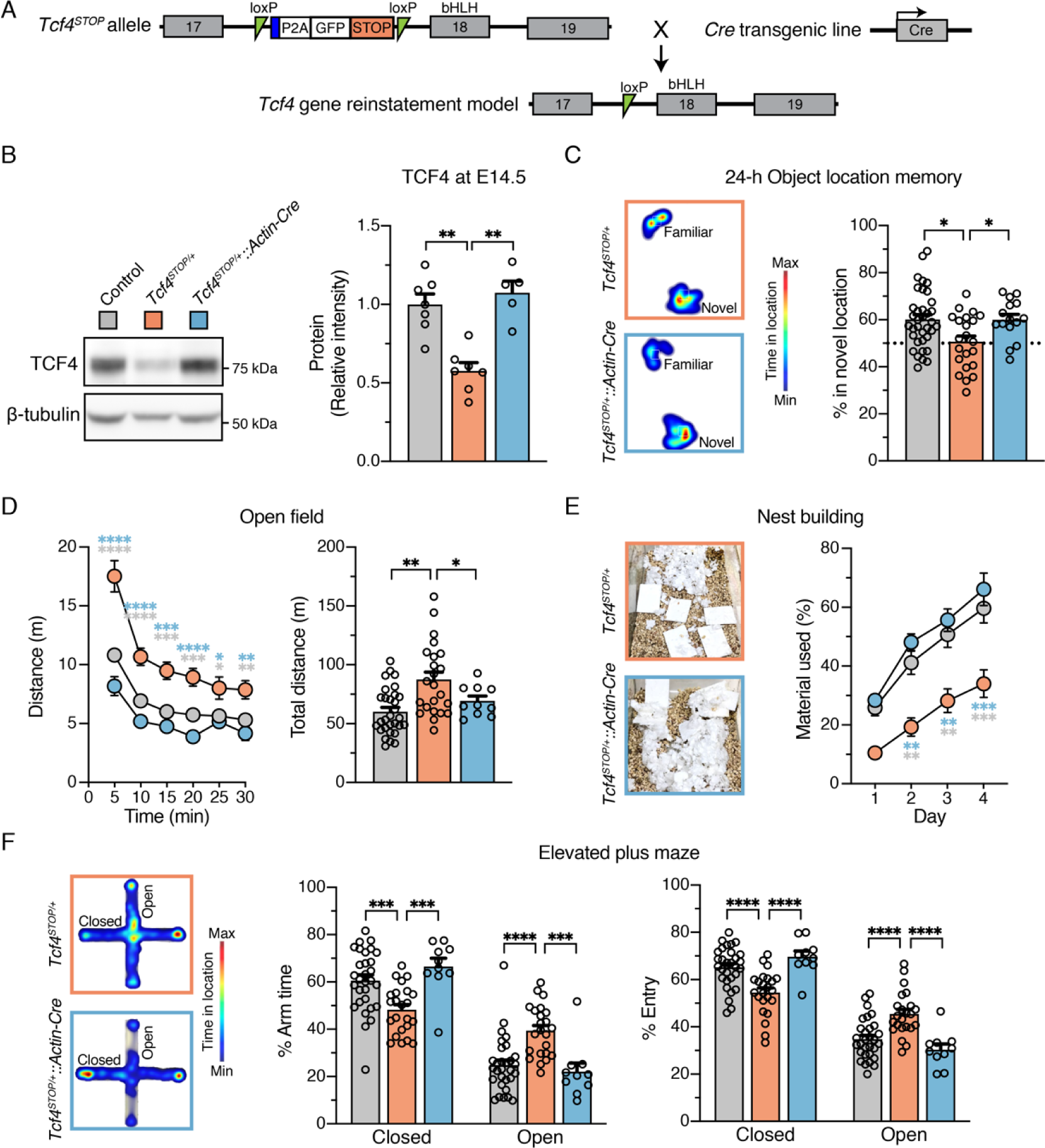
Embryonic, pan-cellular reinstatement of *Tcf4* fully rescues behavioral deficits in a mouse model of Pitt-Hopkins syndrome. (**A**) Schematic depicting a conditional Pitt-Hopkins syndrome mouse model in which expression of the bHLH region of *Tcf4* is prevented by the insertion of a floxed GFP-STOP cassette into intron 17 of *Tcf4* (*Tcf4^STOP/+^*). Crossing *Tcf4^STOP/+^* mice with *Actin-Cre^+/–^* transgenic mice can produce mice with embryonic pan-cellular reinstatement of *Tcf4* expression (*Tcf4^STOP/+^::Actin-Cre*). (**B**) Representative Western blot for TCF4 and b- tubulin loading control protein and quantification of relative intensity of TCF4 protein in embryonic brain lysates from *Tcf4^+/+^* (n=7), *Tcf4^STOP/+^* (n=7), and *Tcf4^STOP/+^::Actin-Cre* (n=5) mice. The data were analyzed by one-way ANOVA followed by Bonferroni’s *post hoc*. (**C**) Left panel: Heatmaps indicate time spent in proximity to one object located in the familiar position and the other object relocated to the novel position. Right panel: Percent time interacting with the novel location object (*Tcf4^+/+^*: n=36, *Tcf4^STOP/+^*: n=22, *Tcf4^STOP/+^::Actin-Cre*: n=15). (**D**) Left panel: Distance traveled per 5 min. Right panel: Total distance traveled for the 30-min testing period (*Tcf4^+/+^*: n=30, *Tcf4^STOP/+^*: n=23, *Tcf4^STOP/+^::Actin-Cre*: n=10). (**E**) Left panel: Representative images of nests built by *Tcf4^STOP/+^* and *Tcf4^STOP/+^::Actin-Cre* mice. Right panel: Percentage of nest material used during the 4-day nest building period (*Tcf4^+/+^*: n=13, *Tcf4^STOP/+^*: n=10, *Tcf4^STOP/+^::Actin-Cre*: n=5). (**F**) Left panel: Heatmaps reveal relative time spent on the elevated plus maze. Right panels: Percent time spent in the closed and open arms and percent of entries made into the closed and open arms (*Tcf4^+/+^*: n=30, *Tcf4^STOP/+^*: n=23, *Tcf4^STOP/+^::Actin-Cre*: n=10). Values are means ± SEM. *p < 0.05, **p < 0.005, ***p < 0.001, ****p < 0.0001.

### *Tcf4* reinstatement in glutamatergic or GABAergic neurons rescues selective behavioral phenotypes in PTHS model mice

Viral-mediated gene delivery can target discrete cell types by promoter choice (Deverman, Ravina, Bankiewicz, Paul, & Sah, 2018), providing a capacity to adjust *Tcf4* expression in a cell type-specific manner. A previous anatomical study shows that TCF4 is present in excitatory and inhibitory neurons of the forebrain (Kim, Berens, Ochandarena, & Philpot, 2020). Moreover, single-cell transcriptomic studies in the neonatal and adult mouse brain indicate that *Tcf4* transcript levels are higher in excitatory and inhibitory neurons than most other cell types (**Figure 2-figure supplement 1**) (Loo et al., 2019; Zeisel et al., 2015). Thus, PTHS-associated pathologies might be effectively treated by preferentially reactivating *Tcf4* expression in excitatory and inhibitory neurons. To explore this possibility, and whether these broad neuronal subclasses contribute to PTHS phenotypes in a modular or cooperative fashion (or both) in the case of *Tcf4* haploinsufficiency, we crossed *Tcf4^STOP/+^* mice to *Nex-Cre^+/-^* or *Gad2-Cre^+/-^* to reactivate *Tcf4* expression selectively in forebrain glutamatergic neurons (*Tcf4^STOP/+^::Nex-Cre* mice) or GABAergic neurons (*Tcf4^STOP/+^::Gad2-Cre* mice) (**Figure 2A**). We then analyzed behavioral outcomes in these mice. In the open field test, we found that *Tcf4^STOP/+^::Nex-Cre* mice exhibited significantly higher activity levels than control mice (*Tcf4^+/+^* and *Tcf4^+/+^::Nex-Cre*) and similar activity levels as *Tcf4^STOP/+^* mice, indicating that embryonic reinstatement of *Tcf4* in forebrain glutamatergic neurons failed to rescue the hyperactivity phenotype (**Figure 2B**). Activity levels in *Tcf4^STOP/+^::Gad2-Cre* mice were statistically indistinguishable from either control mice (*Tcf4^+/+^* and *Tcf4^+/+^::Gad2-Cre)* or *Tcf4^STOP/+^* mice (**Figure 2B**), suggesting that embryonic *Tcf4* reinstatement in GABAergic neurons is also insufficient to fully prevent the hyperactivity phenotype. *Tcf4* reinstatement in glutamatergic neurons improved object location memory, whereas *Tcf4* reinstatement in GABAergic neurons failed to fully prevent location memory impairments (**Figure 2C**). Importantly. lack of improvement in activity level and object location memory in *Tcf4^STOP/+^::Gad2-Cre* mice was reproduced by an independent investigator as part of the same study (**Figure 2-figure supplement 2**). In the elevated plus maze task, we found that *Tcf4^STOP/+^::Nex-Cre* and control mice exhibited increased closed arm activity compared to *Tcf4^STOP/+^* mice, showing that reinstating *Tcf4* in glutamatergic neurons restored the low-anxiety phenotype (**Figure 2D**). In contrast, *Tcf4^STOP/+^::Gad2-Cre* and *Tcf4^STOP/+^* mice exhibited reduced closed arm activity compared to controls (**Figure 2D**), indicating persistence of the reduced anxiety-like phenotype despite reinstatement of *Tcf4* in GABAergic neurons.

**Figure 2.**
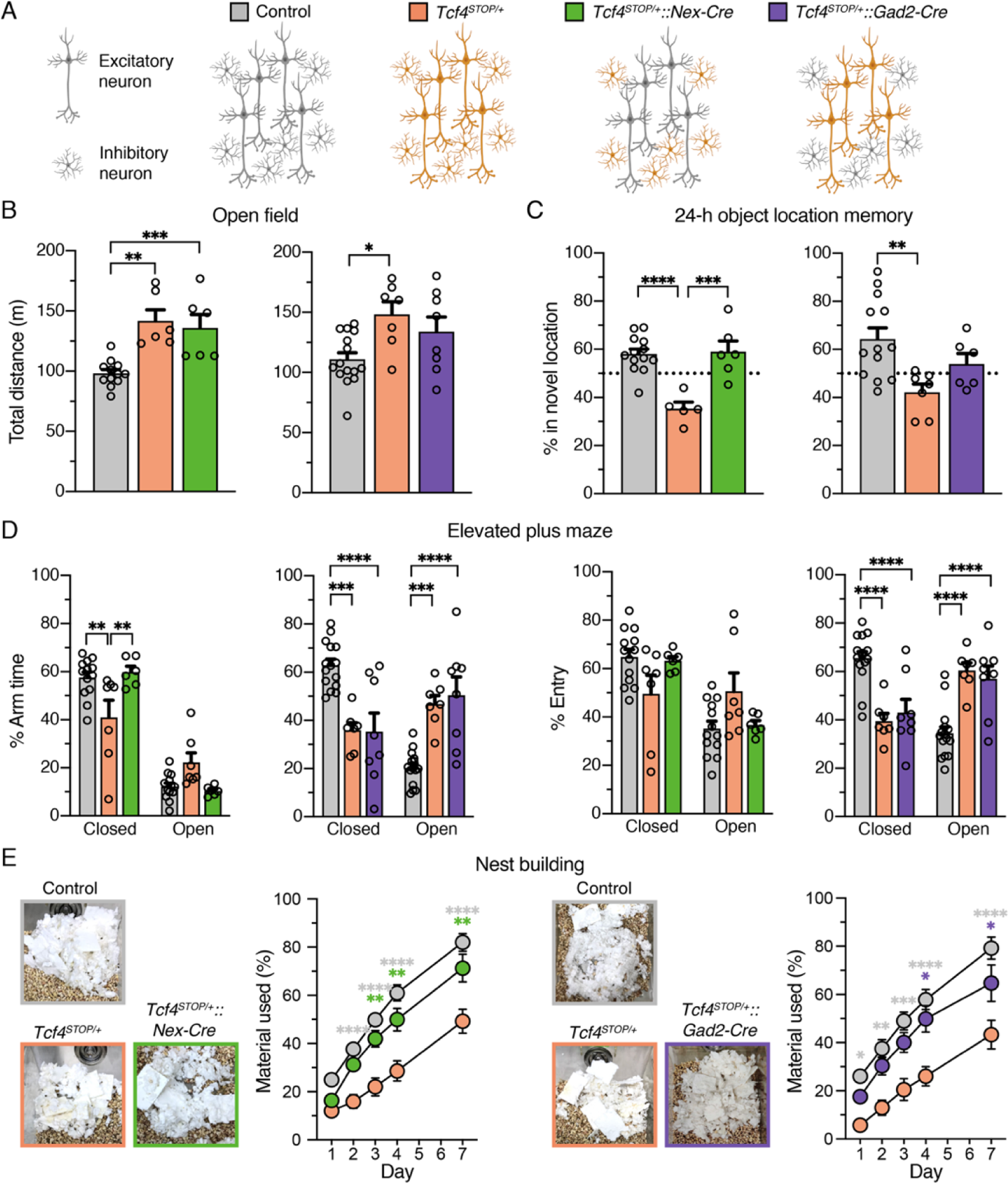
Embryonic reinstatement of *Tcf4* expression in glutamatergic or GABAergic neurons rescues selective behavioral deficits in a mouse model of PTHS. (**A**) Schematic representation of cell type-specific *Tcf4* reinstatement strategy. *Tcf4^STOP/+^::Nex-Cre* or *Tcf4^STOP/+^::Gad2-Cre* mice normalize *Tcf4* expression in glutamatergic or GABAergic neurons, respectively, while controls (*Tcf4^+/+^*, *Nex-Cre^+/–^*, or *Gad2-Cre^+/–^* mice) have normal *Tcf4* expression. (**B**) Total distance traveled for the 30-min testing period (Left panel: *Tcf4^+/+^*: n=12, *Tcf4^STOP/+^*: n=6, *Tcf4^STOP/+^::Nex-Cre*: n=6, and right panel: *Tcf4^+/+^*: n=15, *Tcf4^STOP/+^*: n=7, *Tcf4^STOP/+^::Gad2-Cre*: n=8). (**C**) Percent time interacting with the novel location object (Left panel: *Tcf4^+/+^*: n=13, *Tcf4^STOP/+^*: n=5, *Tcf4^STOP/+^::Nex-Cre*: n=6, and right panel: *Tcf4^+/+^*: n=13, *Tcf4^STOP/+^*: n=7, *Tcf4^STOP/+^::Gad2-Cre*: n=6). (**D**) Percent time spent in closed and open arms and percent entries made into the closed and open arms (Left panel: *Tcf4^+/+^*: n=13, *Tcf4^STOP/+^*: n=7, *Tcf4^STOP/+^::Nex-Cre*: n=6, and right panel: *Tcf4^+/+^*: n=15, *Tcf4^STOP/+^*: n=7, *Tcf4^STOP/+^::Gad2- Cre*: n=8). (**E**) Representative images of nests built by mice and percentage of nest material used during the 7-day nest building period (Left panel: *Tcf4^+/+^*: n=13, *Tcf4^STOP/+^*: n=7, *Tcf4^STOP/+^::Nex-Cre*: n=6, and right panel: *Tcf4^+/+^*: n=15, *Tcf4^STOP/+^*: n=7, *Tcf4^STOP/+^::Gad2-Cre*: n=8). Values are means ± SEM. *p < 0.05, **p < 0.005, ***p < 0.001, ****p < 0.0001.

Finally, we observed that both *Tcf4^STOP/+^::Nex-Cre* and *Tcf4^STOP/+^::Gad2-Cre* mice used a similar amount of nest materials as their respective controls (**Figure 2E**), demonstrating that embryonic reinstatement in either glutamatergic or GABAergic neurons was sufficient to prevent the impaired nest building phenotype in PTHS model mice. Our findings suggest that normalizing *Tcf4* expression from both glutamatergic and GABAergic neurons might be required to fully rescue behavioral phenotypes.

### Neonatal intracerebroventricular administration of PHP.eB-hSyn-Cre produces widespread Cre expression in the brain during early postnatal development

We aimed to establish the extent to which postnatal reinstatement of *Tcf4* could prevent behavioral deficits in PTHS model mice. We conducted experiments designed to mimic an eventual viral-mediated gene therapy for PTHS in an idealistic manner with respect to reintroducing wildtype *Tcf4* isoforms and expression levels. To this end, we packaged a Cre transgene cassette into a recombinant AAV9-derived PHP.eB vector and bilaterally delivered this viral vector to the cerebral ventricles of neonates (**Figure 3A**). We expressed the Cre cassette under control of the human synapsin promoter (hSyn) for selective expression in neurons (Nieuwenhuis et al., 2021) (**Figure 3-figure supplement 1**). We initially examined expression of the Cre cassette as a proxy for the temporal and spatial biodistribution of *Tcf4* reinstatement following intracerebroventricular (ICV) injection of 1 µl of ∼3.2 x 10^10^ vg AAV9/PHP.eB-hSyn-Cre on postnatal day 1 (P1) (**Figure 3A**). We failed to detect significant Cre signals from relatively medial sagittal sections of P4 and P7 mouse brain, despite being able to observe a local distribution of Cre-expressing cells in the brain regions near the lateral ventricle injection site (**Figure 3B-C** and **Figure 3-figure supplement 1B-C**). Cre mRNA and protein distribution across the forebrain remained sparse until at least P10 (**Figure 3B-C** and **Figure 3-figure supplement 1D**). But, Cre was visibly and more broadly expressed in the hippocampus and throughout the cortical layers by P17 (**Figure 3B-C** and **Figure 3-figure supplement 1E**). The biodistribution of Cre protein at P60 was widespread in the brain, with particularly prominent expression in the forebrain compared to subcortical regions (**Figure 3D-F**), which is similar to patterns of endogenous TCF4 distribution (compare **Figure 3D** and Figure 3-figure supplement 1B).

**Figure 3.**
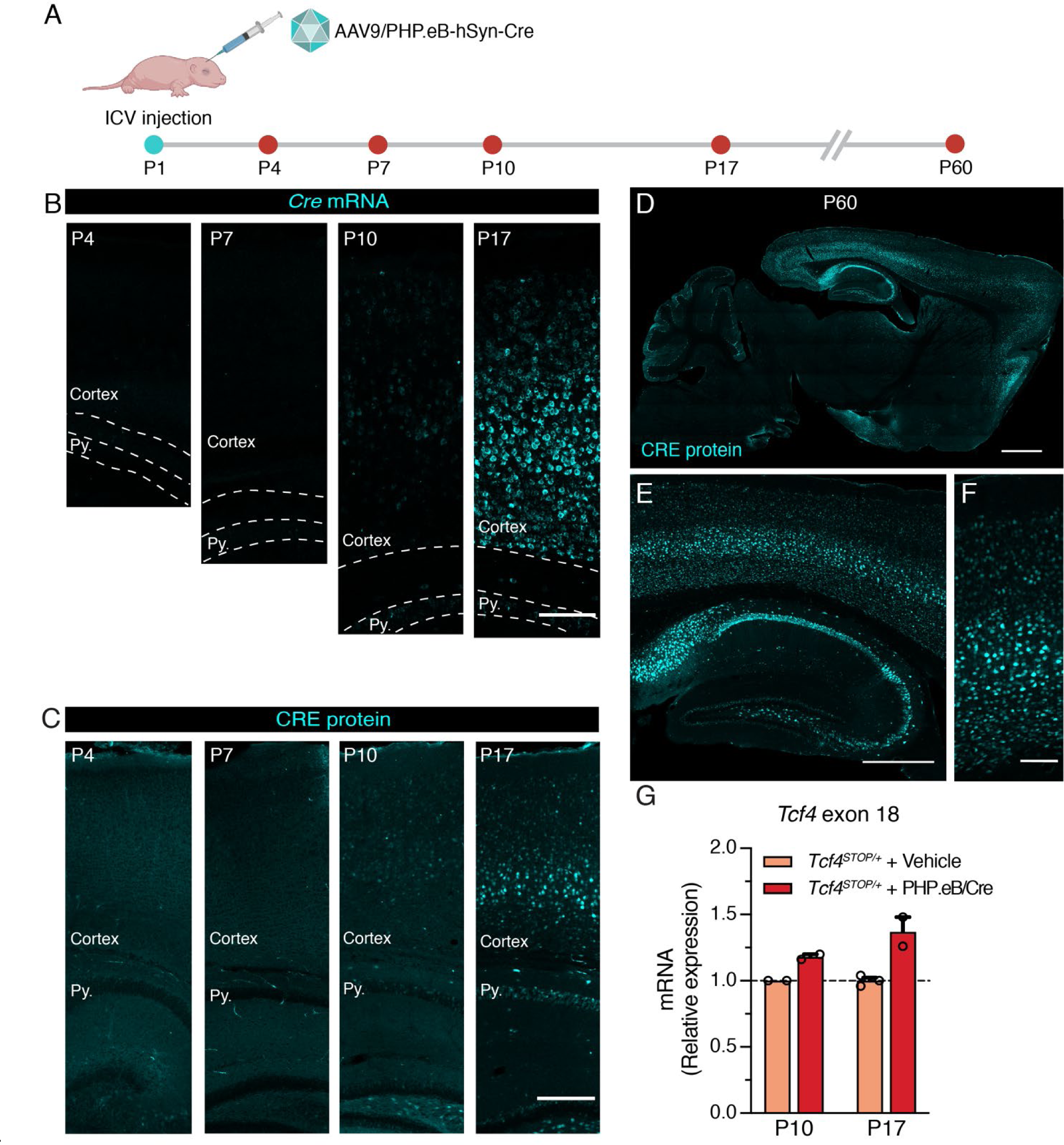
Neonatal ICV delivery of PHP.eB/Cre yields Cre expression by approximately P10-P17. (**A**) A timeline of experiment to evaluate timing of Cre biodistribution following intracerebroventricular (ICV) injection of 1 µl of 3.2 x 10^13^ vg/ml AAV9/PHP.eB-hSyn-Cre to P1 mice. (**B**) *In situ* hybridization for *Cre* mRNA, and (**C**) immunofluorescence staining for CRE protein in the cortex and hippocampus of P4, P7, P10, and P17 wildtype mice neonatally treated with PHP.eB/Cre. Py. = Stratum pyramidale. Scale bars = 100 µm (B) and 250 µm (C). (**D-F**) CRE immunofluorescence staining in sagittal section of P60 wildtype mouse brain. Scale bars = 1 mm (D), 500 mm (E), and 100 mm (F). (**G**) Relative *Tcf4* transcript levels detected in the brains of P10 *Tcf4^STOP/+^* mice treated with vehicle (n=2) or PHP.eB/Cre (n=2) and P17 *Tcf4^STOP/+^* mice treated with vehicle (n=3) or PHP.eB/Cre (n=2).

To test whether Cre expression coincides with *Tcf4* reinstatement in *Tcf4^STOP/+^* mice, we quantified full-length *Tcf4* mRNA transcripts upon delivery of PHP.eB/Cre to *Tcf4^STOP/+^* neonates. The RT-qPCR results confirmed increased relative expression of *Tcf4* transcripts from P10 and P17 *Tcf4^STOP/+^* brains treated with PHP.eB/Cre (**Figure 3G**, *Tcf4^STOP/+^* + Vehicle at P10: 1.0 ± 0.0, n = 2; *Tcf4^STOP/+^* + PHP.eB/Cre at P10: 1.18 ± 0.03, n = 2, *Tcf4^STOP/+^* + Vehicle at P19: 1.0 ± 0.04, n = 3; *Tcf4^STOP/+^* + PHP.eB/Cre at P19: 1.37 ± 0.16, n = 2). Taken together, these observations confirm neonatal ICV injection to be a viable route of delivery to examine the behavioral consequences of postnatal *Tcf4* reinstatement in a subset of neurons.

### Postnatal reinstatement of *Tcf4* expression ameliorates behavioral phenotypes in PTHS model mice

We analyzed the behavioral performance of adult (P60 - P110) *Tcf4^+/+^* and *Tcf4^STOP/+^* mice after delivering vehicle or PHP.eB/Cre at P1 (**Figure 4A**). Similar to vehicle- or virally-treated *Tcf4^+/+^* mice, *Tcf4^STOP/+^* mice treated with PHP.eB/Cre exhibited normal activity levels in the open field test, whereas vehicle-treated *Tcf4^STOP/+^* mice were hyperactive (**Figure 4B**). In addition to normalizing activity levels, PHP.eB/Cre treatment also fully rescued long-term memory performance in *Tcf4^STOP/+^* mice compared to vehicle-treated *Tcf4^STOP/+^* mice (**Figure 4C**). In the elevated plus maze, PHP.eB/Cre-treated *Tcf4^STOP/+^* mice spent relatively more time in the closed arms than the open arms, similar to vehicle- and PHP.eB/Cre-treated *Tcf4^+/+^* mice. In contrast, vehicle-treated *Tcf4^STOP/+^* mice spent similar time in the open and closed arms as a sign of their abnormally low anxiety levels (**Figure 4D**). Lastly, we found that PHP.eB/Cre treatment supported progressive improvement of nest building behavior in *Tcf4^STOP/+^* mice. Specifically, although both *Tcf4^STOP/+^* groups appeared to be similarly lacking in the execution of this behavior at baseline, over the course of 1 week, PHP.eB/Cre-treated *Tcf4^STOP/+^* mice proved capable of incorporating nest materials to a similar degree as vehicle- or virally-treated *Tcf4^+/+^* mice (**Figure 4E**). While PHP.eB/Cre-mediated postnatal reinstatement of *Tcf4* partially or fully recovered performance on a variety of behavioral phenotypes in PTHS model mice, we found that small body and brain sizes were not corrected (**Figure 4F-G**). Earlier intervention and/or improved pan-cellular biodistribution might be necessary to restore anatomical integrity, although this is evidently not a prerequisite for behavioral recovery. Collectively, these data demonstrate the potential for virally-mediated postnatal normalization of *Tcf4* expression to broadly rescue behavioral phenotypes.

**Figure 4.**
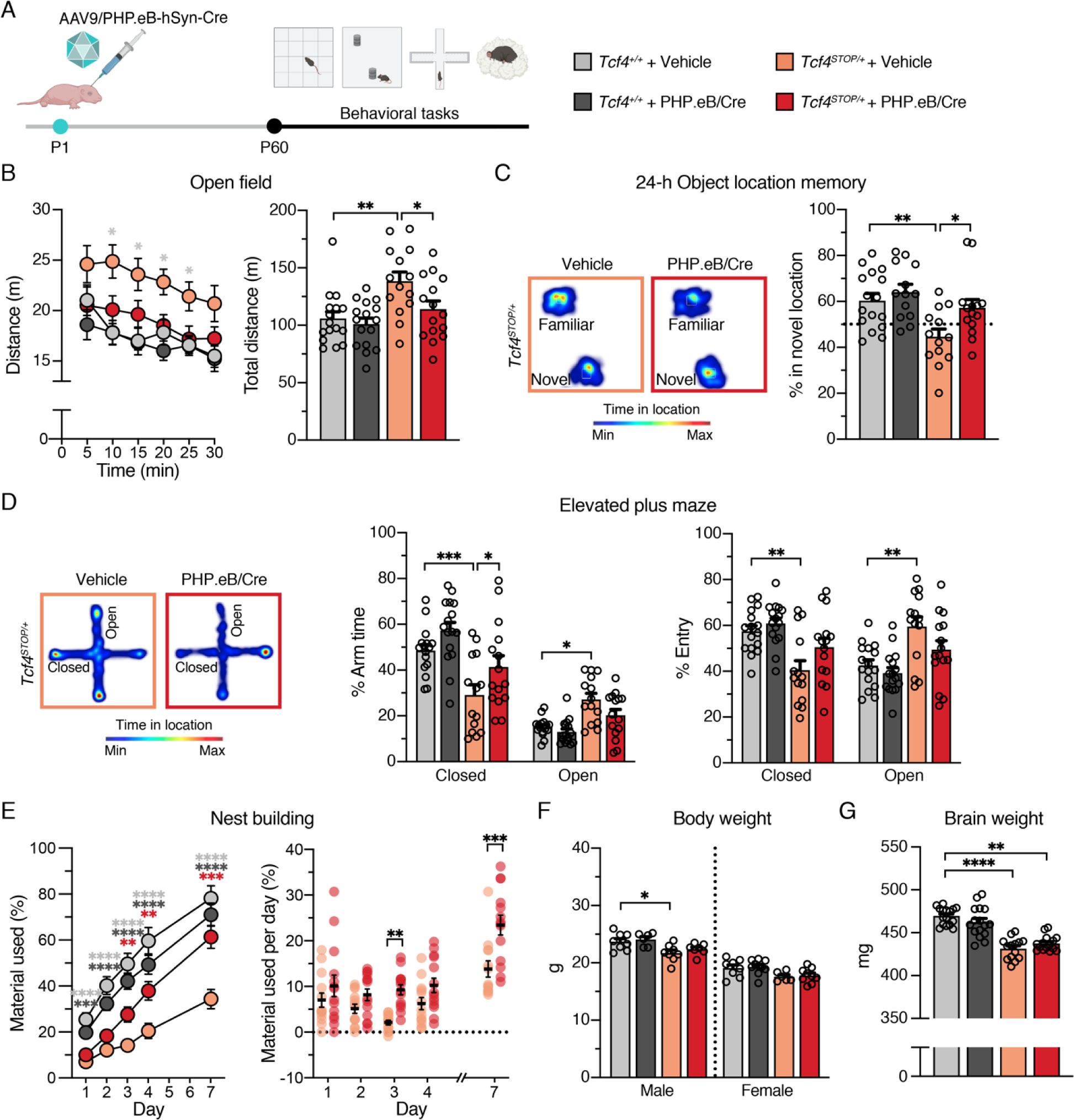
Neonatal ICV injection of PHP.eB/Cre improves behavioral phenotypes in *Tcf4^STOP/+^* mice. (**A**) Experimental timeline for evaluation of behavioral phenotypes in *Tcf4^+/+^* and *Tcf4^STOP/+^* mice treated with vehicle or PHP.eB/Cre. (**B**) Left panel: Distance traveled per 5 min. Right panel: Total distance traveled for the 30-min testing period. (**C**) Left panel: Heatmaps indicate time spent in proximity to one object located in the familiar position and the other object relocated to a novel position. Right panel: Percent time interacting with the novel location object. (**D**) Left panel: Heatmaps reveal time spent in elevated plus maze. Right panels: Percent time spent in the closed and open arms and percent entries made into the closed and open arms. (**E**) Left panel: Percentage of nest material used during the 7-day nest building period. Right panel: Percentage of nest material used per day. (**F**) Body weight analysis of P65-69 male and female mice. (**G**) Adult brain weight analysis. Values are means ± SEM. *p < 0.05, **p < 0.005, ***p < 0.001, ****p < 0.0001.

### Postnatal *Tcf4* reinstatement partially corrects local field potential abnormalities in PTHS model mice

Several clinical observations have reported electroencephalographic (EEG) abnormalities, such as altered slow waves, in individuals with PTHS (Amiel et al., 2007; Peippo et al., 2006; Takano, Lyons, Moyes, Jones, & Schwartz, 2010), yet these phenotypes have not been examined in PTHS model mice. Here we performed local field potential (LFP) recordings in *Tcf4^STOP/+^* mice, which provide an accurate indication of local neuronal activity (Buzsaki, Anastassiou, & Koch, 2012). We implanted recording electrodes in the hippocampus, a site of high *Tcf4* expression (Kim et al., 2020), and recorded LFP from freely moving mice (**Figure 5A** and **Figure 5-figure supplement 1A-B**). We observed a trend for reduced total LFP power in *Tcf4^STOP/+^* mice, but the total power between *Tcf4^+/+^* and *Tcf4^STOP/+^* mice was not statistically distinguishable (**Figure 5-figure supplement 1C-D**). Significant decreases in *Tcf4^STOP/+^* LFP power were evinced in the theta (5-8 Hz) band (**Figure 5-figure supplement 1C, E**). A moderate but consistent decrease in power likely underlie this phenotype as per follow-up spectrogram analyses (**Figure 5-figure supplement 1F**). Having established that *Tcf4* haploinsufficiency resulted in LFP abnormalities in mice, we sought to determine whether LFP power could be normalized by postnatal *Tcf4* reinstatement. Upon analyzing LFP recordings in vehicle- and virus-treated groups (**Figure 5B- C**), we found total LFP power to be significantly reduced in vehicle-treated *Tcf4^STOP/+^* mice compared to vehicle- and virally-treated *Tcf4^+/+^* mice, but partially normalized in PHP.eB/Cre- treated *Tcf4^STOP/+^* mice (**Figure 5D**). This effect appeared to be largely driven by the normalization of theta band activity (**Figure 5E**), which was evident across one-minute recording epochs (**Figure 5F**). Collectively, these data define hippocampal LFP deficits in PTHS model mice and demonstrate their amenability to normalization by postnatal reinstatement of *Tcf4*.

**Figure 5.**
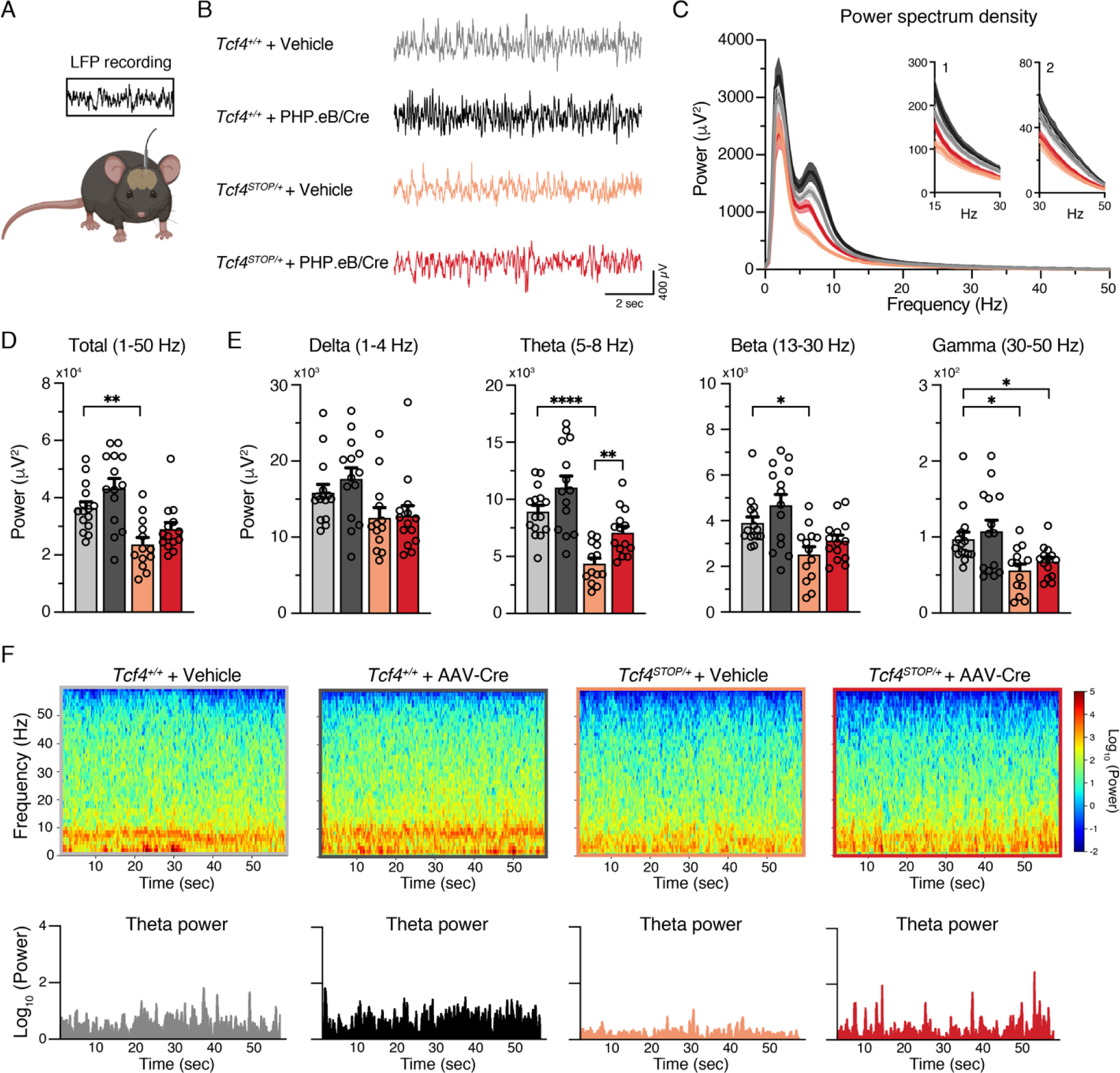
Neonatal ICV injection of PHP.eB/Cre partially rescues LFP spectral power in *Tcf4^STOP/+^* mice. (**A**) Schematic of LFP recording from the hippocampus of a freely moving mouse. (**B**) Representative examples of LFP in each experimental group. (**C**) Power spectrum density of hippocampal LFP analyzed from *Tcf4^+/+^* and *Tcf4^STOP/+^* mice treated with vehicle or PHP.eB/Cre. Inset 1 spans from 15 to 30 Hz and inset 2 spans from 30 to 50 Hz on x- axis. (**D**) LFP power analyses of frequency bands ranging from 1 to 50 Hz, (**E**) delta (1-4 Hz), theta (5-8 Hz), beta (13-30 Hz), and gamma (30-50 Hz) bands (*Tcf4^+/+^* + vehicle: n=15, *Tcf4^+/+^* + PHP.eB/Cre: n=14, *Tcf4^STOP/+^* + vehicle: n=13, and *Tcf4^STOP/+^* + PHP.eB/Cre: n=14). (**F**) Top panel: Spectrograms in single LFP sessions of representative experimental groups. Bottom panel: Representative theta power extracted from spectrogram in the top panel. Values are means ± SEM. *p < 0.05, **p < 0.005, ****p < 0.0001.

### Postmortem evaluation of *Cre* biodistribution and expression of *Tcf4* and TCF4-regulated genes

After completing behavioral and LFP experiments, we performed *in situ* hybridization (ISH) to characterize *Cre* distribution and RT-qPCR to examine effectiveness of PHP.eB/Cre treatment on expression levels of *Tcf4* and TCF4-regulated genes. ISH revealed that *Cre* mRNA was still robustly detected in 6-month-old mice, subsequent to viral delivery of the *Cre* transgene at P1 (**Figure 6A-D**). In the cortex and olfactory bulb, *Cre* was observed in most cells, but certainly not all, throughout the layers (**Figure 6C-D**). Similarly, although we found most pyramidal cells to express *Cre* in the hippocampus, non-expressing cells were evident, especially in the dentate gyrus (**Figure 6B**). This observation suggests that reinstating *Tcf4* in a subset of neurons can provide therapeutic benefit. To ensure the relative uniformity of *Cre* transduction among treated *Tcf4^STOP/+^* mice, we analyzed *Cre* fluorescence in the neocortex and pyramidal cell layer of CA1. Levels proved generally consistent, with the exception of one mouse whose neocortex and CA1 *Cre* fluorescence was ∼3-4 times higher than the group median (**Figure 6E**).

**Figure 6.**
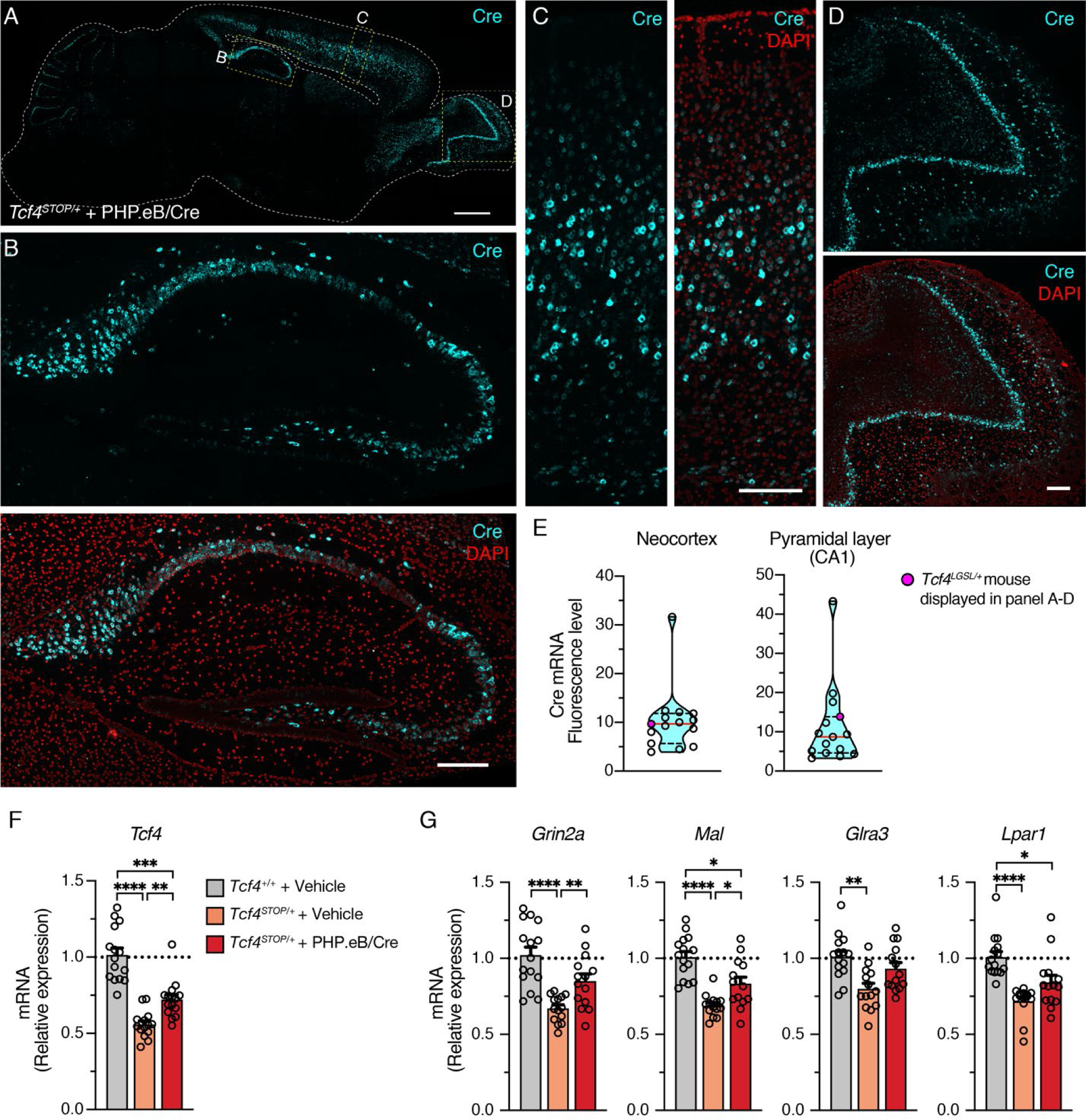
Widespread *Cre* expression of the forebrain leads to partial upregulation of *Tcf4* and partial recovery of selected TCF4-regulated gene expression. (**A**) Representative image of *in situ* hybridization for *Cre* mRNA in sagittal section of six-month-old *Tcf4^STOP/+^* mouse that was treated at P1 with PHP.eB/Cre. Scale bar = 1 mm (**B-D**) Higher magnification images of boxed regions in panel A. Scale bars = 200 mm. (**E**) *Cre* mRNA fluorescence levels of neocortex and CA1 pyramidal cell layer analyzed from individual *Tcf4^STOP/+^* + PHP.eB/Cre mice. The red line and black dotted lines of the violin plot represent median and interquartile ranges of the data, respectively. (**F**) Relative *Tcf4* mRNA expression of the forebrain from vehicle-treated *Tcf4^+/+^* (n=15) and *Tcf4^STOP/+^* (n=14) mice and PHP.eB/Cre-treated *Tcf4^STOP/+^* (n=15) mice. *Tcf4* mRNA expression levels of PHP.eB/Cre-treated *Tcf4^STOP/+^* mice are relatively higher than vehicle-treated *Tcf4^STOP/+^* mice. (**G**) Relative mRNA expressions of selected TCF4-regulated genes. Transcript levels of *Grin2a* (encoding for NMDA receptor subunit epsilon-1), *Mal* (encoding for myelin and lymphocyte protein), *Glra3* (encoding for glycine receptor subunit alpha-3), and *Lpar1* (encoding for lysophosphatidic acid receptor 1) were partially upregulated by *Tcf4* reinstatement. Values are means ± SEM. ***p < 0.001, ****p < 0.0001.

We next analyzed *Tcf4* mRNA expression from the forebrains of the PHP.eB/Cre-treated *Tcf4^STOP/+^* mice. On average, *Tcf4* mRNA expression was approximately 1.3-fold higher in virally-treated versus vehicle-treated *Tcf4^STOP/+^* forebrains at P150-P200 (**Figure 6F**, *Tcf4^+/+^* + Vehicle: 1.0 ± 0.18, n = 15; *Tcf4^STOP/+^* + Vehicle: 0.56 ± 0.09, n = 14; *Tcf4^STOP/+^* + PHP.eB/Cre: 0.72 ± 0.12, n = 15). Recent RNA-sequencing studies revealed genes whose expression levels were altered by heterozygous *Tcf4* disruption (Kennedy et al., 2016; Phan et al., 2020). To analyze the impact of PHP.eB/Cre treatment on the expression of TCF4-regulated genes, we examined expression levels of the following genes: *Grin2a*, *Mal*, *Glra3*, and *Lpar1*, whose expression has been shown to be dysregulated by *Tcf4* haploinsufficiency (Kennedy et al., 2016; Phan et al., 2020). Each was noticeably downregulated in vehicle-treated *Tcf4^STOP/+^* mice, but at least partially normalized by PHP.eB/Cre treatment (**Figure 6G**). Our postmortem analyses confirmed that neonatal PHP.eB/Cre treatment in *Tcf4^STOP/+^* mice partially normalized expression levels of both *Tcf4* and TCF4-targeted genes. Thus, modestly increased *Tcf4* expression was sufficient to rescue behavioral phenotypes in *Tcf4^STOP/+^* mice, suggesting that treatment approaches based on *TCF4* normalization may benefit from a relatively wide therapeutic window.

## Discussion

The genetic mechanism of PTHS suggests a therapeutic opportunity; loss-of-function in one *TCF4* copy is sufficient to cause PTHS, so conversely, restoring TCF4 function could treat PTHS. In this proof-of-concept study, genetic reinstatement of *Tcf4* during early postnatal development corrected multiple behavioral phenotypes in PTHS model mice, including hyperactivity, reduced anxiety-like behavior, memory deficit, and abnormal innate behavior. Furthermore, early postnatal *Tcf4* reinstatement corrected altered local field potential activity and, at the molecular level, TCF4-regulated gene expression changes. Our results suggest that postnatal genetic therapies to compensate for loss-of-function of TCF4 can offer an effective treatment for PTHS.

One of the key parameters for genetic normalization strategies is the age at time of intervention. We likely accomplished widespread *Tcf4* normalization throughout the brain by P10-P17 (**Figure 3**), which is roughly equivalent to the first two years of human life (S. Wang, Lai, Deng, & Song, 2020). Individuals with PTHS have global developmental delay, often presenting itself in the first year of life (Zollino et al., 2019). Thus, our study indicates that genetic normalization approaches could provide a viable early life treatment opportunity. Future investigation is needed to address the extent to which juvenile and adult *Tcf4* reinstatement can correct behavioral phenotypes, as the therapeutic window could be broader than what we describe in this study.

While it is necessary to intervene within a developmental therapeutic window, eventual ASO- or AAV-mediated genetic therapies will also need to produce an appropriate biodistribution to be efficacious. Previous studies have shown that *Tcf4* expression levels are particularly high in the forebrain (Kim et al., 2020). Accordingly, our proof-of-concept study employed a strategy to reinstate *Tcf4* expression more prominently in the forebrain than in subcortical regions (**Figure 3** and **6**). Notably, intrathecal delivery of ASOs produces a biodistribution favoring the forebrain (Mazur et al., 2019), remarkably similar to the endogenous *Tcf4* expression pattern (Kim et al., 2020). Given our ability to recover behavioral phenotypes, future efforts to test the feasibility of ASOs or gene therapy should target *Tcf4* reinstatement in the forebrain to provide therapeutic benefit.

Another important consideration for eventual genetic therapies is which cell types should be targeted to rescue behavioral phenotypes. The therapeutic potential of genetic rescue in specific cell types had not been evaluated due to the lack of conditional models for *Tcf4* reinstatement. Our conditional restoration model provides a powerful tool in that we can establish the cellular and behavioral impacts of cell type-specific *Tcf4* restoration, which will ultimately inform therapeutic development for PTHS. Because *Tcf4* is particularly expressed at high levels in glutamatergic and GABAergic neurons during embryonic development (Jung et al., 2018) and throughout the postnatal period (Kim et al., 2020) (**Figure 2-figure supplement 1**), we initially aimed to embryonically reinstate TCF4 function in these broad neuronal classes to establish their relative contribution to behavioral rescue. Reactivating *Tcf4* expression in excitatory pyramidal neurons, dentate gyrus mossy cells, and granule cells within the dorsal telencephalon, starting from E11.5, improved memory, anxiety phenotype, and innate behavior, while reactivating *Tcf4* expression in almost all GABAergic neurons throughout the brain at ∼E13.5 rescued only abnormal innate behavior (Goebbels et al., 2006; Taniguchi et al., 2011) (**Figure 2**). Our data suggest that TCF4 in both neuronal types contribute to behavioral outcomes, and therapeutic strategies should thus target the normalization of *Tcf4* expression in both glutamatergic and GABAergic neurons. Accordingly, we demonstrated, at the proof-of-concept level, a potential for phenotypic rescue when TCF4 function was postnatally restored in all neuronal types (**Figure 4-5**). However, we cannot rule out the possibility of gaining phenotypic rescue by reinstating *Tcf4* from other cell types such as oligodendrocytes in which the loss of TCF4 has been shown to contribute to PTHS pathophysiology (Phan et al., 2020).

A *TCF4* normalization treatment strategy has an advantage in that it addresses the core genetic defect in PTHS, and therefore should restore transcriptional targets of TCF4. *Tcf4* haploinsufficiency altered the expression of genes that are involved in synaptic plasticity and neuronal excitability, such as *Grin2a* and *Glra3*, and neuronal development, such as *Mal* and *Lpar1* (Kennedy et al., 2016; Phan et al., 2020). Our data show that upregulating *Tcf4* levels postnatally can correct the expression of these TCF4-regulated genes (**Figure 6**). The effect of *Tcf4* normalization on downstream genes might help to guide future preclinical studies. For example, therapeutic agent choice and their dosing could be optimized by testing behavioral recovery and by measuring expression of TCF4-regulated genes, such as those validated in this study.

Our study has several important limitations. First, we normalized *Tcf4* expression only in neurons. *Tcf4* is expressed in nearly all neurons, astrocytes, and oligodendrocytes (Jung et al., 2018; Kim et al., 2020). Ideally, *Tcf4* should be reinstated in both neuronal and non-neuronal cells to accomplish maximum therapeutic outcomes. Our preliminary data indicated that injecting AAV containing a broadly active promoter, CAG, reinstated *Tcf4* in all cell types, but induced abnormal glial activation, which is reminiscent of toxicity previously reported with CAG vectors (Xiong et al., 2019), and severe weight loss (Data not shown). To avoid toxicity in our experimental paradigm while still achieving efficient transduction, we employed the neuron-selective promoter (hSyn) in our viral construct. Nonetheless, our data provide compelling evidence that reinstating *Tcf4* only in neuronal cells is sufficient to reverse behavioral and LFP phenotypes in PTHS model mice. Second, our study does not inform therapeutic threshold that must be achieved by genetic normalization approaches. TCF4 is a dosage-sensitive protein: too little expression causes neurodevelopmental disorders, and too much expression appears to be linked to schizophrenia (Brennand et al., 2011; Brzozka, Radyushkin, Wichert, Ehrenreich, & Rossner, 2010; M. Forrest et al., 2012; Quednow, Brzozka, & Rossner, 2014; Sepp et al., 2012; Wirgenes et al., 2012). Our conditional model allowed us to establish the best-case treatment scenario by reinstating *Tcf4* to wildtype levels. Future proof-of-concept preclinical studies to upregulate *Tcf4* through ASOs or gene therapy approaches in PTHS model mice must take considerable care to recapitulate optimal levels of *Tcf4* expression. Third, our study does not guide appropriate isoforms to deliver to the brain through AAV-mediated gene therapy. The *TCF4* gene produces at least 18 isoforms, which may have cell type- and developmental-specific expression patterns in the brain (Sepp, Kannike, Eesmaa, Urb, & Timmusk, 2011).

Characterizing endogenous isoform expression in the human brain will be critical to guide design of a viral vector that produces appropriate *TCF4* isoform expression.

To our knowledge, the present study is the first investigation to show that normalizing *Tcf4* expression during early postnatal development can improve behavioral outcomes in a mouse model of PTHS. Furthermore, our studies provide insights into target cell types and biodistribution that must be achieved for therapeutic recovery in mice, which guides the rational design of genetic normalization approaches such as AAV-mediated gene therapy, ASOs, or small molecules. In sum, our findings suggest parameters for which genetic therapies can provide substantial therapeutic benefit for individuals with PTHS.

## Materials and Methods

### Study design

Wildtype females were crossed to *Tcf4^STOP/+^* males to generate wildtype and *Tcf4^STOP/+^* mice (PTHS model mice). *Tcf4^STOP/+^* females were crossed to *Cre* transgenic males to conditionally reinstate *Tcf4* expression in a Cre-dependent manner (**Figure 1 and 2**). Neonatal (P1-2) *Tcf4^STOP/+^* and *Tcf4^+/+^* mice were randomly assigned to treatment with vehicle or AAV9/PHP.eB-hSyn-Cre at a dose of 3.2 x 10^10^ vector genomes (vg) delivered bilaterally to the cerebral ventricles. All injected mice performed a battery of behavioral tests beginning 2 months of age, spanning a period of 6-7 weeks, in the following order: Open field, object location memory, elevated plus maze, and nest building (**Figure 4**). All the treated mice underwent electrode implantation surgery 2 weeks after the last behavioral test and recovered from the surgery for at least 7 days. Most treated mice with intact electrode headcaps were subjected to LFP recording. LFP data were acquired for 3 days, 1 hour each day (**Figure 5**). Upon the completion of LFP recording, mice were sacrificed for *in situ* hybridization (ISH) and qPCR analyses. Half a brain was used for ISH staining, and the other half was used for qPCR experiment (**Figure 6**). All behavioral and LFP data were from two biological replicates, and all behavioral experiments were performed only once. All investigators who conducted experiments and analyzed data were blinded to genotype and treatment until completion of the study.

### Mice

The generation of *Tcf4^STOP/+^* knock-in mice has been previously described (Kim et al., 2020). Mice carrying *loxP-GFP-STOP-loxP* allele were maintained on a congenic C57BL/6J background. The female *Tcf4^STOP/+^* mice were mated with heterozygous males from one of three *Cre*-expressing lines: *Nex-Cre^+/-^* (Goebbels et al., 2006), which Klaus-Armin Nave generously provided, *Gad2-Cre^+/-^* (RRID:IMSR_JAX:010802), or *Actin-Cre^+/-^* (RRID:IMSR_JAX:019099). All mice were maintained on a 12:12 light-dark cycle with *ad libitum* access to food and water. We used male and female littermates at equivalent genotypic ratios. All research procedures using mice were approved by the Institutional Animal Care and Use Committee at the University of North Carolina at Chapel Hill and conformed to National Institutes of Health guidelines.

### Calculation of Tcf4 expression from public single-cell sequencing data

Single-cell transcriptomic data from the neonatal mouse cortex (Loo et al., 2019) and the adult mouse nervous system (Zeisel et al., 2015) were obtained from GEO accession GSE123335 and from http://mousebrain.org/downloads.html, respectively. For the neonatal cortex data, the mean and standard error of *Tcf4* expression values were computed across all cells of a given annotated cell type in R and plotted using ggplot2. For adult data, we focused just on cell types annotated as deriving from cortex, amygdala, dentate gyrus, hippocampus, olfactory bulb, cerebellum, and striatum. We then grouped similar cell types into broader classifications. For example, all clusters annotated as glutamatergic (GLU) were renamed as “Excitatory neuron”. We then computed the mean and standard error of *Tcf4* expression for each broader cell type within each brain region in R and plotted using ggplot2. All code to reproduce the plots for Figure 2-figure supplement 1 is provided at https://github.com/jeremymsimon/Kim_TCF4.

### Adeno-associated viral vector production

To produce AAV9/PHP.eB capsids, a PEI triple transfection protocol was first performed. Then the product was grown under serum-free conditions and purified through three rounds of CsCl density gradient centrifugation. Purified product was exchanged into storage buffer containing 1 x phosphate-buffered saline (PBS), 5% D-Sorbitol, and 350mM NaCl. Virus titers (GC/ml) were determined by qPCR targeting the AAV inverted terminal repeats. A codon-optimized *Cre* cDNA was packaged into AAV9/PHP.eB capsids.

### AAV delivery

P1-2 mouse pups were cryo-anesthetized on ice for about 3 minutes, then transferred to a chilled stage equipped with a fiber optic light source for transillumination of the lateral ventricles. A 10 μl syringe fitted with a 32-gauge, 0.4-inch-long sterile syringe needle (7803-04, Hamilton) was used to bilaterally deliver 0.5 μl of AAV9/PHP.eB-hSyn-Cre or vehicle (PBS supplemented with 5% D-Sorbitol and additional 212 mM NaCl) to the ventricles. The addition of Fast Green dye (1 mg/mL) to the virus solution visualized injection area. Following injection, pups were warmed on an isothermal heating pad with home-cage nesting material before being returned to their home cages.

### Behavioral testing and analyses

Object location memory: Mice were habituated to an open box, containing a visual cue on one side, without objects for 5 minutes each day for 3 days. In the following day, mice were trained with two identical objects for 10 minutes. After 24 hours, mice were placed in the box where one of the objects was relocated to a novel position for 5 minutes. Video was recorded during each period. Interaction time of a mouse with each object was measured by Ethovision XT 15.0 program (Noldus). A percentage of the exploration time with the object in a novel position (% in novel location) was calculated as follows: (time exploring novel location)/(time exploring novel location + familiar location) * 100. If total exploration time was less than 2 seconds, these mice were excluded from the dataset.

Open field (**Figure 1D**): Mice were given a 30-min trial in an open-field chamber (41 x 41 x 30 cm) that was crossed by a grid of photobeams (VereMax system, Accuscan Instruments). Counts were taken of the number of photobeams broken during the trial in 5-min intervals. Total distance traveled was measured over the course of the 30-min trial.

Open field (**Figure 2B** and **4B**): Mice were given a 30-min trial in an open-field chamber (40 x 40 x 30 cm). Mouse movements were recorded with a video camera, and the total distance traveled was measured by Ethovision XT 15.0 program.

Elevated plus maze: The elevated plus maze was constructed to have two open arms and two closed arms; all arms are 20 cm in length and 8 cm in width. The maze was elevated 50 cm above the floor. Mice were placed on the center section and allowed to explore the maze for 5 minutes. Mouse movements on the maze were recorded with a video camera. Activity levels (time and entry) in open or closed arms were measured by Ethovision XT 15.0 program.

Nest building: Mice were single-housed for a period of 3 days before the start of the assay. On day 1, 10-11 g of compressed extra-thick blot filter paper (1703966, Bio-Rad), cut into 8 evenly sized rectangles, was placed in a cage. In each day, for 4 consecutive days (**Figure 1E**), the amount of paper not incorporated into a nest was weighed. For **Figure 2E** and **4E**, additional measurement of nest material was recorded 72 hours after collecting data for 4 consecutive days.

### Surgery and in vivo LFP recording

Mice were anaesthetized by inhalation of 1-1.5% isoflurane (Piramal) in pure O2 during surgery, with 0.25% bupivacaine injected under the scalp for local analgesia and meloxicam (10 mg/kg) subcutaneously administered. Stainless steel bipolar recording electrodes (P1 Technologies) were implanted in the hippocampus (coordinates from bregma: AP = -1.82 mm; ML= 1.5 mm; and DV = -1.2 mm), and ground electrodes were fastened to a stainless-steel screw positioned on the skull above the cerebellum. Dental cement was applied to secure electrode positions. Mice recovered for at least 7 days prior to LFP recording. A tethered system with a commutator (P1 Technologies) was used for recordings, while mice freely moved in their home cages. LFP recordings were amplified (1000x) using single-channel amplifiers (Grass Technologies), sampled at a rate of 1000 Hz, and filtered at 0.3 Hz high-pass and 100 Hz low-pass filters. All electrical data were digitized with a CED Micro1401 digital acquisition unit (Cambridge Electronic Design Ltd.).

### LFP analysis

Data acquired in Spike2 software (Cambridge Electronic Design Ltd.) were read into Python and further processed with a butter bandpass filter from 1-100 Hz to focus on frequencies of our interest. Frequency bands were defined as delta 1-4 Hz, theta 5-8 Hz, beta 13-30 Hz, and gamma 30-50 Hz. Spectral power was analyzed using the Welch’s Method, where the power spectral density is estimated by dividing the data into overlapping segments. Sample size (“n”) represents the number of mice. For each mouse, we selected the longest continuous period with no movement artifacts for analysis. We averaged processed data obtained across three days. We wrote custom Python scripts to analyze LFP data.

### Tissue preparation

Mice were anesthetized with sodium pentobarbital (60 mg/kg, intraperitoneal injection) before transcranial perfusion with 25 ml of PBS immediately followed by phosphate-buffered 4% paraformaldehyde (pH 7.4). Brains were postfixed overnight at 4°C before 24-hour incubations in PBS with 30% sucrose. Brains were sectioned coronally or sagittaly at 40 μm using a freezing sliding microtome (Thermo Scientific). Sections were stored at -20°C in a cryo-preservative solution (45% PBS, 30% ethylene glycol, and 25% glycerol by volume).

### Immunohistochemistry

For immunofluorescent staining, sections were rinsed several times with PBS (pH = 7.3) and PBS containing 0.2% Triton X-100 (PBST) before blocking with 5% normal goat serum in PBST (NGST) for 1 hour at room temperature (RT). Sections were then incubated with primary antibodies diluted in NGST at 4°C for 48 hours. Sections were rinsed several times with PBST and then incubated with secondary antibodies for 1 hour at RT. In all experiments, 4’,6- diamidino-2-phenylindole (DAPI; Invitrogen D1306) was added during the secondary antibody incubation at a concentration of 700 ng/ml. Primary antibodies used included 1:1000 mouse anti-Cre (Millipore, MAB3120), 1:1000 guinea pig anti-NeuN (Millipore, ABN90P), and 1:1000 rabbit anti-GFAP (Dako, Z0334). The following secondary antibodies from Invitrogen (Carlsbad, CA) were used at 1:1000 dilution: goat anti-mouse Alexa 647 (A21240), goat anti-guinea pig Alexa 594 (A11076), and goat anti-rabbit Alexa 568 (A11011).

For chromogenic staining, sections were rinsed several times with PBS, and endogenous peroxidases were quenched by incubating for 5 mins in 1.0% H2O2 in MeOH, followed by PBS rinsing. Sections were washed with PBST several times. Then sections were blocked with 5% NGST for 1 hour at RT. Blocked sections were incubated in primary antibody, rabbit anti-GFP (NB600-308) diluted 1:1000 in NGST for 24 hours at 4°C. After incubation in primary antibodies, sections were rinsed several times in PBST and incubated for 1 hour at RT in biotinylated goat anti-rabbit secondary antibodies (Vector BA-1000, Burlingame, CA) diluted 1:500 in NGST. Sections were then rinsed in PBST prior to tertiary amplification for 1 hour with the ABC elite avidin-biotin-peroxidase system (Vector PK-7100). Further rinsing with PBST preceded a 3-minute incubation at RT in 3’3’-diaminobenzidine (DAB) chromogenic substrate (0.02% DAB and 0.01% H2O2 in PBST) to visualize immune complexes amplified by avidin-biotin-peroxidase.

### In situ hybridization

Brains were extracted and frozen in dry ice. Sections were taken at a thickness of 16 μm. Staining procedure was completed to manufacturer’s specifications. RNAscope Fluorescent Multiplex Assay (Advanced Cell Diagnostics), designed to visualize multiple cellular RNA targets in fresh frozen tissues (F. Wang et al., 2012), was used to detect Cre (Cat No. 423321-C3) in mouse sections.

### Imaging

Images of brain sections stained by using fluorophore-conjugated secondary antibodies were obtained with Zeiss LSM 710 Confocal Microscope, equipped with ZEN imaging software (Zeiss) and a Nikon Ti2 Eclipse Color and Widefield Microscope (Nikon). Images compared within the same figures were taken using identical imaging parameters. Images within figure panels went through identical modification for brightness and contrast by using Fiji Image J software. Figures were prepared using Adobe Illustrator software (Adobe Systems).

### RT-qPCR

The neocortical and hippocampal hemispheres were rapidly dissected, snap-frozen with dry ice-ethanol bath, and stored at -80°C. Total RNAs were extracted using the RNeasy Mini Kit (74106, Qiagen), and reverse transcribed via qScript cDNA SuperMix (101414-106, QuantaBio). The resulting cDNAs constituted the input, and RT-qPCR was performed in a QuantStudio Real-Time PCR system using SYBR green master mix (A25742, Thermofisher). The specificity of the amplification products was verified by melting curve analysis. All RT-qPCRs were conducted in technical triplicates, and the results were averaged for each sample, normalized to *Actin* expression, and analyzed using the comparative ΔΔCt method. The triplicates are valid only when the standard deviation is smaller than 0.25. The following primers were used in the RT-qPCRs: m*Tcf4* (forward: 5’-GGGAGGAAGAGAAGGTGT-3’, reverse: 5’- CATCTGTCCCATGTGATTCGC-3’), *Grin2a* (forward: 5’- TTCATGATCCAGGAGGAGTTTG-3’, reverse: 5’-AATCGGAAAGGCGGAGAATAG-3’), *Mal* (forward: 5’-CTGGCCACCATCTCAATGT-3’, reverse: 5’- TGGACCACGTAGATCAGAGT-3’), *Glra3* (forward: 5’-GGGCATCACCACTGTACTTA-3’, reverse: 5’-CCGCCATCCAAATGTCAATAG-3’), *Npy2r* (forward: 5’- GAAGTGAAAGTGGAGCCCTATG-3’, reverse: 5’- ATCTTGCTCTCCAGGTGGTA-3’), *Npar1* (forward: 5’- CCCTCTACAGTGACTCCTACTT-3’, reverse: 5’- GCCAAAGATGTGAGCGTAGA-3’), and *Actin* (forward: 5’- GGCACCACACCTTCTACAATG-3’, reverse: 5’-GGGGTGTTGAAGGTCTCAAAC-3’).

### Western blot

Embryonic day 14.5 brains were sonicated on ice using radioimmunoprecipitation assay buffer [50 mM Tris-HCl pH 8.0, 150 mM NaCl, 1% NP-40, 0.5% Na-deoxycholate, 0.5% SDS, and protease inhibitor cocktail (P8340, Sigma)]. Tissue homogenates were cleared by centrifugation at 4°C. Protein concentrations were determined by bicinchoninic acid assay (23225, Thermo Scientific). 20 μg of each sample was separated in 8% sodium dodecyl sulfate–polyacrylamide gel electrophoresis and transferred to polyvinylidene fluoride membranes (45-004-110, Fisher Scientific) in ice-cold transfer buffer (25 mM Tris-base, 192 mM glycine, and 20% MeOH).

Membranes were blocked in Odyssey Blocking Buffer (927-40100, Li-COR Biosciences) for 1 hour and then blotted with primary antibodies overnight at 4°C and then subsequently, with secondary antibodies for 1 hour at RT. The following primary antibodies were used: Mouse anti- TCF4 (1:1000; sc-393407, Santa Cruz Biotechnology) and rabbit anti-β-Tubulin (1:5000; ab6046, Abcam). The following secondary antibodies were used: Horseradish peroxidase-conjugated anti-mouse and anti-rabbit antibodies (1:5000; 31430 and 31460, Thermo Fisher).

Chemiluminescence reaction was performed using Clarity Western ECL Substrate (1705061, Bio-Rad), which was imaged by an Amersham Imager 680 (GE Healthcare). The signal was quantified using the ImageJ software.

### Statistics

Welch’s one-way analysis of variance (ANOVA) followed by Dunnett’s *post hoc* test was performed for brain and body weight, object location memory task, open field (total distance), LFP power analysis, and qPCR results (**Figure 1**, **5**, and **6**). One-way ANOVA followed by Bonferroni’s *post hoc* was carried out for Western blot and qPCR analyses, body and brain weight, object location memory task, and open field (total distance) (**Figure 1, 2,** and **4**). Two-way ANOVA followed by Tukey’s *post hoc* was conducted for elevated plus maze, open field (distance for every 5-min), and nest building (% material used). Two-way ANOVA followed by Bonferroni’s *post hoc* was used to analyze % material used per day for nest building assay (**Figure 4**). All values are expressed as means ± standard error of the mean (SEM). Asterisks indicate *P* values: \**P* < 0.05, \*\**P* < 0.005, \*\*\**P* < 0.001, \*\*\*\**P* < 0.0001. GraphPad Prism 9.1.1 software (GraphPad Software) was used for all statistical analyses.

## Supplementary Materials

Figure 1-figure supplement 1. Body and brain weight analysis.

Figure 2-figure supplement 1. Single-cell RNA sequencing reveals cell type-specific *Tcf4* expression in the neonatal and adult mouse brain.

Figure 2-figure supplement 2. Behavioral outcomes of *Tcf4^STOP/+^::Gad2-Cre* mice were independently replicated.

Figure 3-figure supplement 1. Cre immunofluorescence staining in sagittal sections of P4, P7, P10, and P17 mice.

Figure 5-figure supplement 1. *Tcf4* haploinsufficiency alters LFP spectral power in the theta band.

## Acknowledgments

We thank Matthew C. Judson for critical readings of the manuscript, Dale Cowley at the UNC Animal Models Core for designing and generating the new mouse model, Klaus-Armin Nave for providing *Nex-Cre* mice, and Viktoriya Nikolova for training on behavioral tasks.

---

**Figure 1-figure supplement 1.**
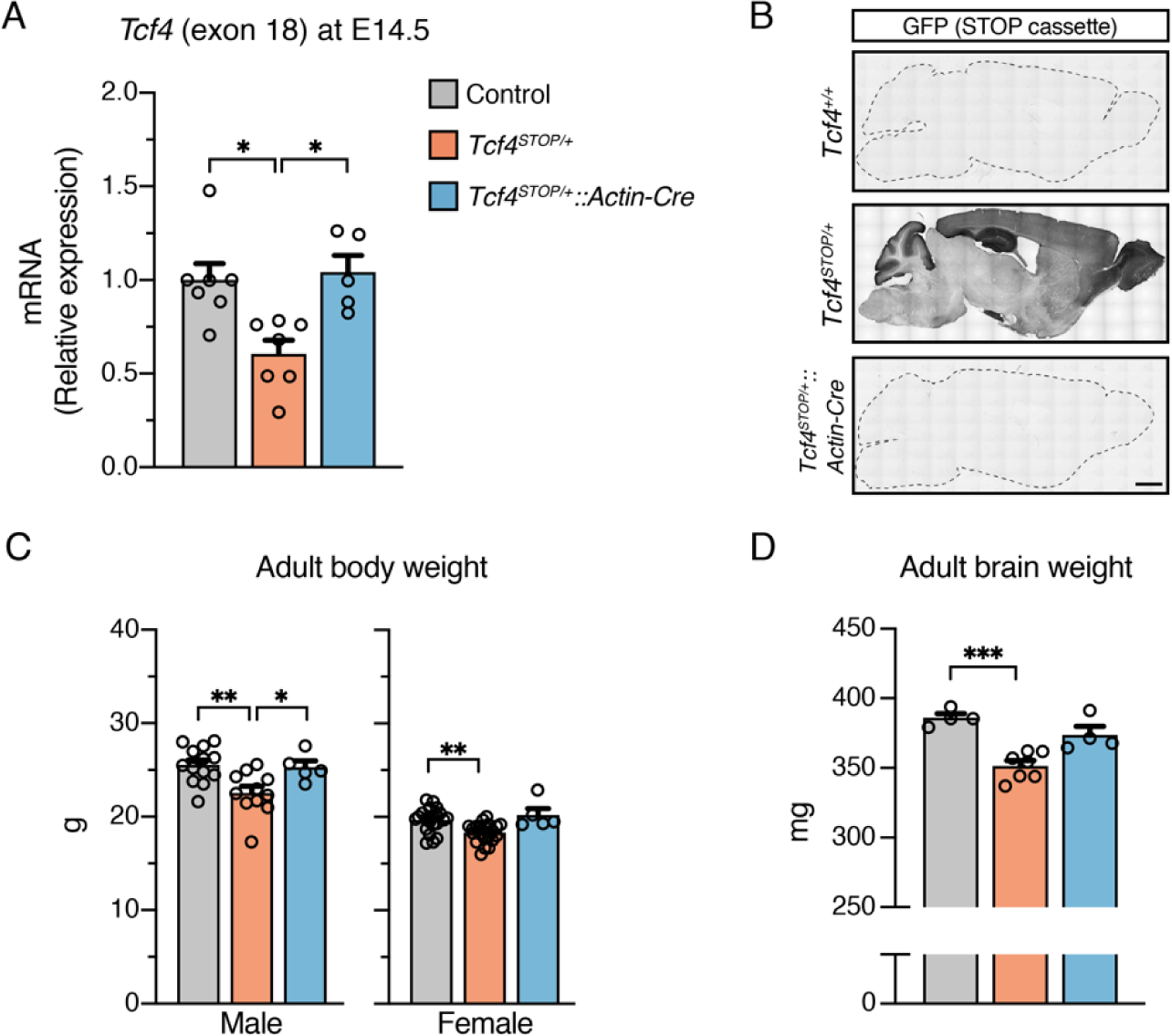
Body and brain weight analysis. **(A)** Relative *Tcf4* mRNA expression in embryonic brain lysates from *Tcf4^+/+^* (n=7), *Tcf4^STOP/+^* (n=7), and *Tcf4^STOP/+^::Actin-Cre* (n=5). **(B)** DAB immunostaining of GFP (indicating presence of the STOP cassette) in sagittal brain sections of adult *Tcf4^+/+^*, *Tcf4^STOP/+^*, and *Tcf4^STOP/+^::Actin-Cre* mice. Scale bar = 1 mm. **(C)** Adult male (*Tcf4^+/+^*: n=13, *Tcf4^STOP/+^*: n=11, *Tcf4^STOP/+^::Actin-Cre*: n=5) and female (*Tcf4^+/+^*: n=22, *Tcf4^STOP/+^*: n=21, *Tcf4^STOP/+^::Actin-Cre*: n=5)body weights of each genotypic group. **(D)** Adult brain weight measured from dissected brains (*Tcf4^+/+^*: n=4, *Tcf4^STOP/+^*: n=7, *Tcf4^STOP/+^::Actin-Cre*: n=4). Values are means ± SEM. *p < 0.05, **p < 0.005, ***p < 0.001.

**Figure 2-figure supplement 1.**
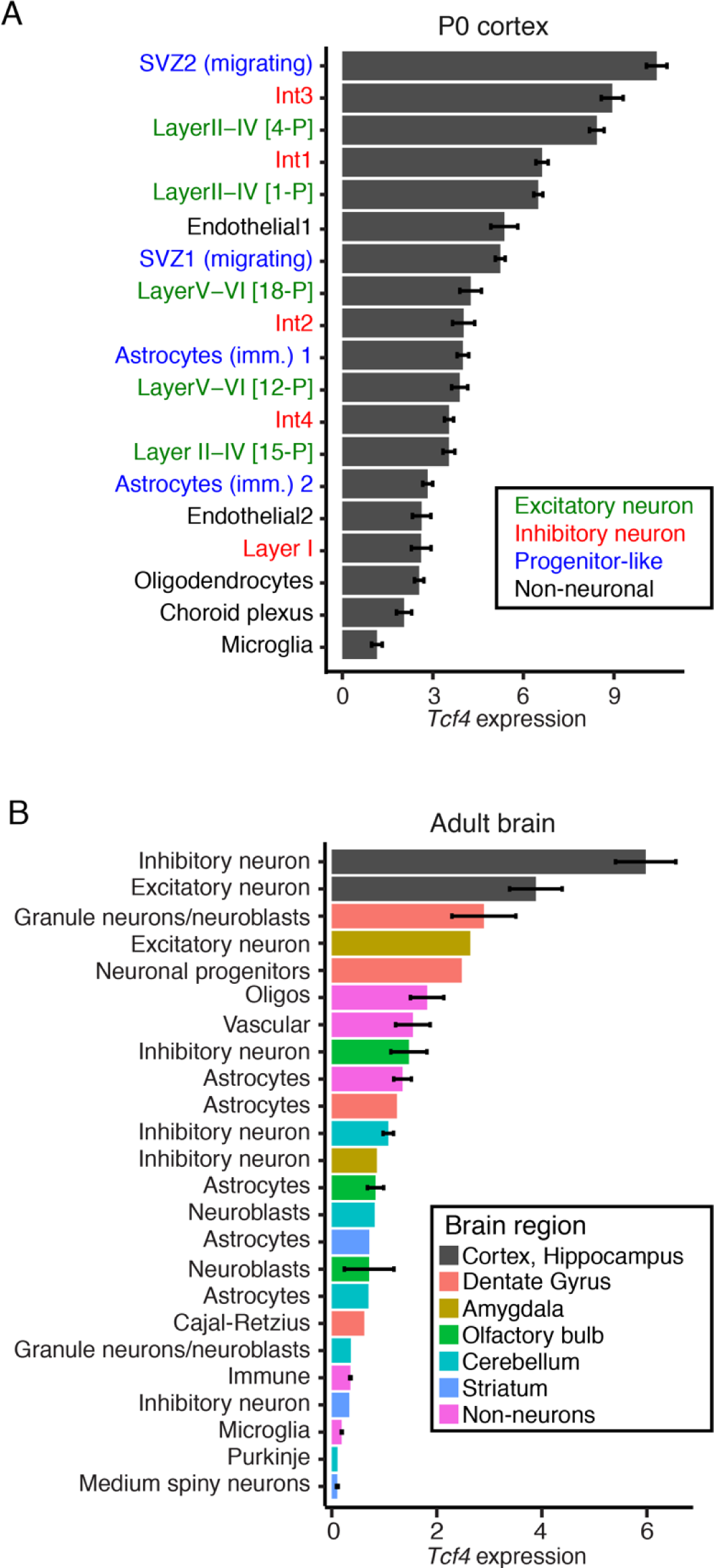
Single-cell RNA sequencing reveals cell type-specific *Tcf4* expression in the neonatal and adult mouse brain. **(A)** Average expression of *Tcf4* in 19 cell types identified in the P0 cortex [data analyzed from reference (Loo et al., 2019)]. Cell type names are colored by principal cell classifications. **(B)** Average expression of *Tcf4* in cell types identified in the adult mouse brain [data analyzed from reference (Zeisel et al., 2015)]. Bars are colored by brain region, and clusters were aggregated into principal cell types. Values are means ± SEM.

**Figure 2-figure supplement 2.**
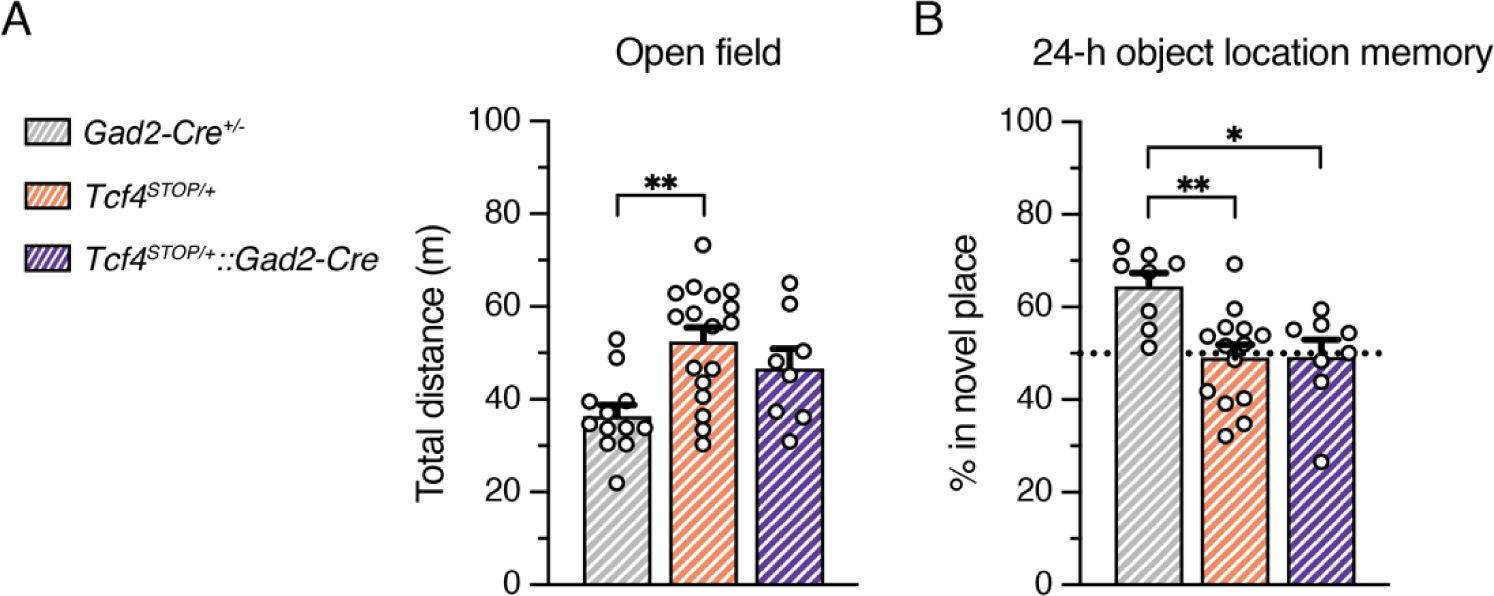
Behavioral outcomes of *Tcf4^STOP/+^::Gad2-Cre* mice were independently replicated. **(A)** Total distance traveled for the 30-min testing period (*Tcf4^+/+^*: n=12, *Tcf4^STOP/+^*: n=17, *Tcf4^STOP/+^::Gad2-Cre*: n=8). **(B)** Percent time interacting with the novel location object. Reinstating *Tcf4* from GABAergic neurons did not rescue hyperactivity and memory function deficit (*Tcf4^+/+^*: n=8, *Tcf4^STOP/+^*: n=14, *Tcf4^STOP/+^::Gad2-Cre*: n=8). Values are means ± SEM. *p < 0.05, **p < 0.005. These experiments were performed at Bates College in the lab of Dr. Andrew Kennedy.

**Figure 3-figure supplement 1.**
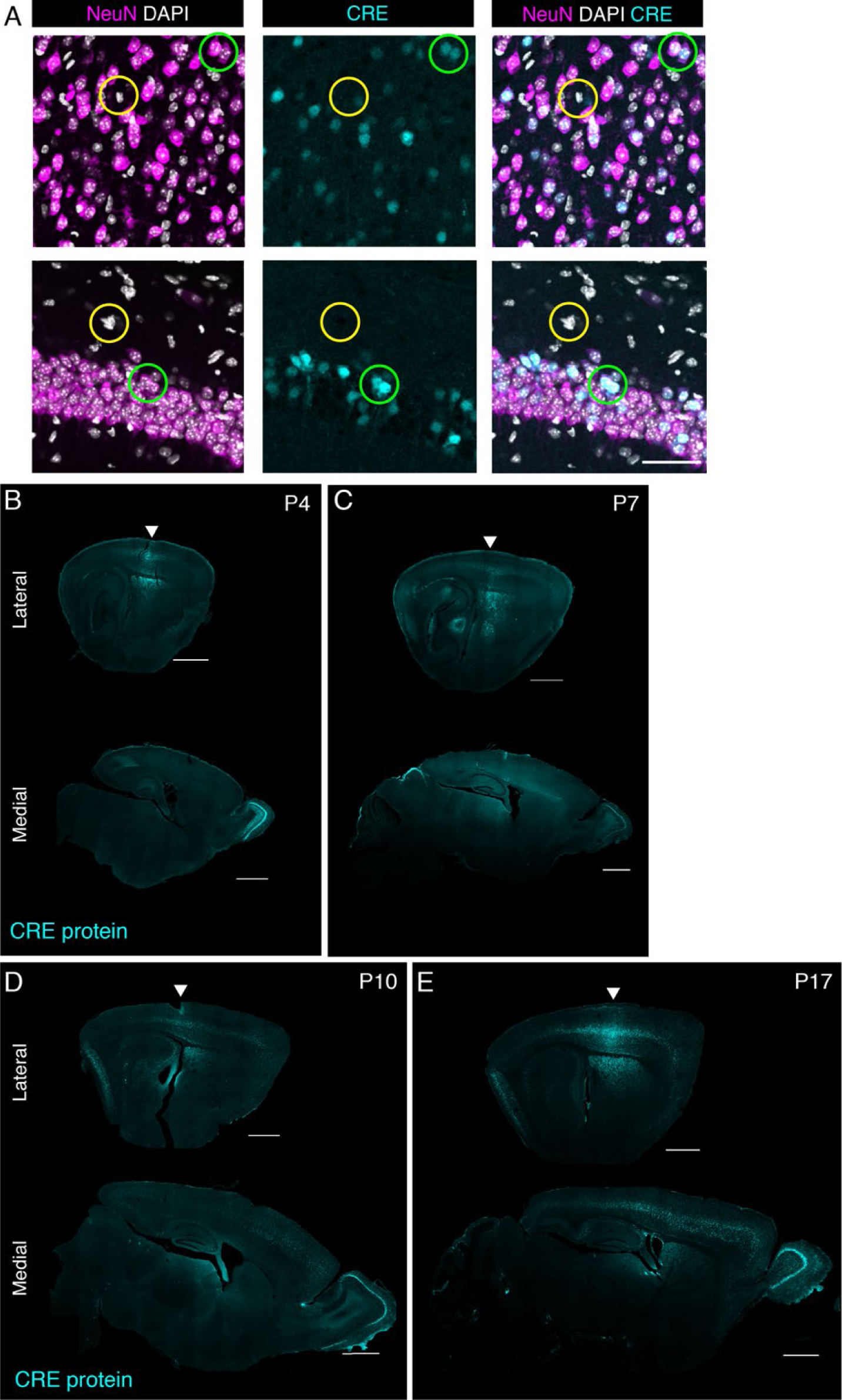
Cre immunofluorescence staining in sagittal sections of P4, P7, P10, and P17 mice. **(A)** Dual immunostaining of neuronal marker (NeuN) and Cre in P17 brain of a mouse treated with PHP.eB/Cre. CRE protein was detected in NeuN-positive cells (green circle), but absent in NeuN-negative cells (yellow circle). Scale bars = 100 µm. **(B-E)** CRE protein spatial pattern in lateral (top row) and medial (bottom row) sagittal sections at different postnatal time points. Arrows indicate the brain area close to the injection site. Scale bars = 1 mm.

**Figure 5-figure supplement 1.**
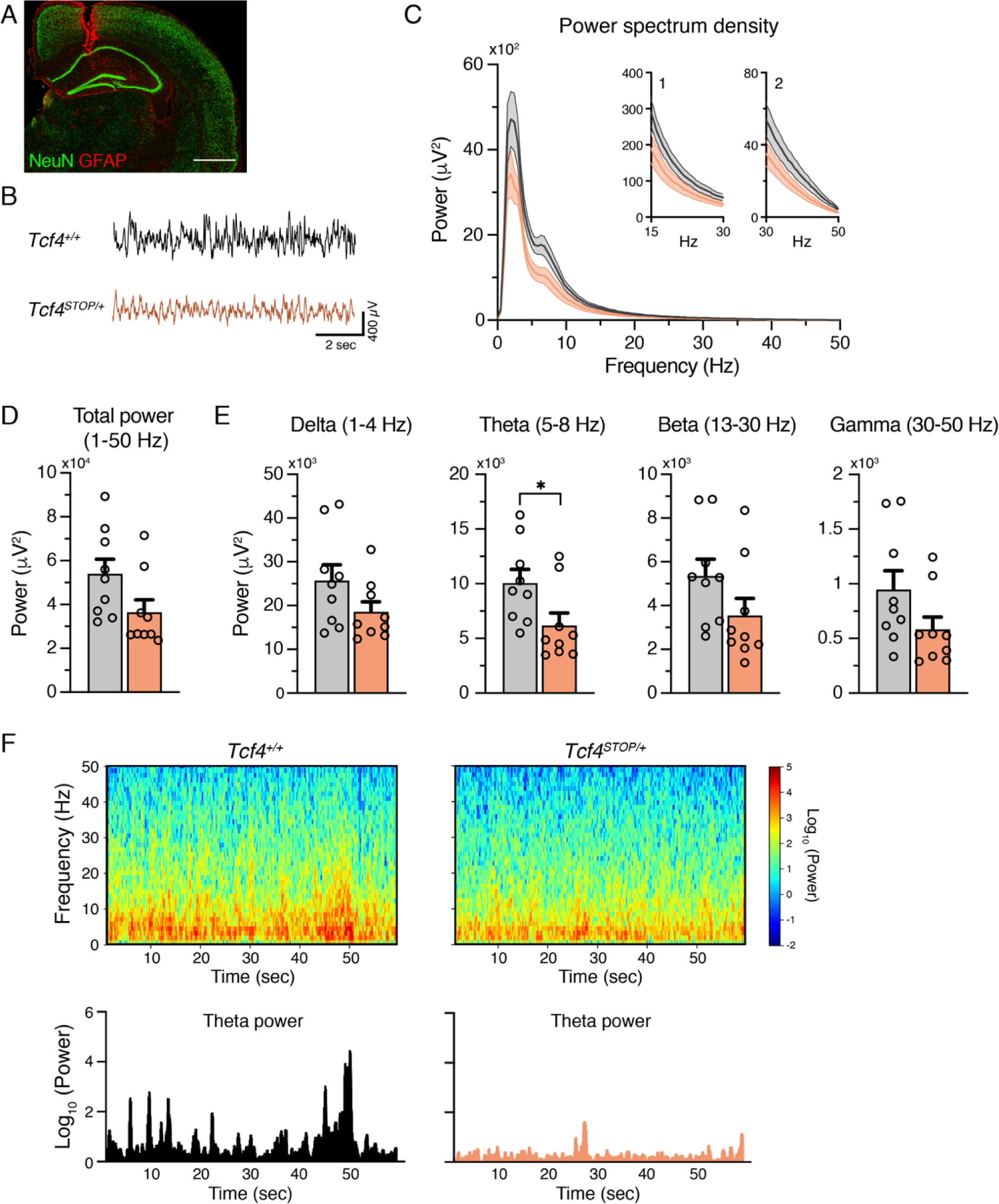
*Tcf4* haploinsufficiency alters LFP spectral power in the theta band. **(A)** Example electrode location (red, GFAP; green, NeuN). Scale bar = 100 mm. **(B)** Representative examples of LFP in *Tcf4^+/+^* and *Tcf4^STOP/+^* mice. **(C)** Power spectrum density of hippocampal LFP analyzed from *Tcf4^+/+^* and *Tcf4^STOP/+^* mice. Inset 1 spans from 15 to 30 Hz and inset 2 spans from 30 to 50 Hz on x-axis. **(D)** LFP power analyses of frequency bands ranging from 1 to 50 Hz, **(E)** delta (1-4 Hz), theta (5-8 Hz), beta (13-30 Hz), and gamma (30-50 Hz) bands (*Tcf4^+/+^*: n=9 and *Tcf4^STOP/+^*: n=9). **(F)** Top panel: Spectrograms in a single LFP session of representative experimental groups. Bottom panel: Representative theta power extracted from spectrogram in the top panel. Values are means ± SEM. *p < 0.05.

**Table 1.**
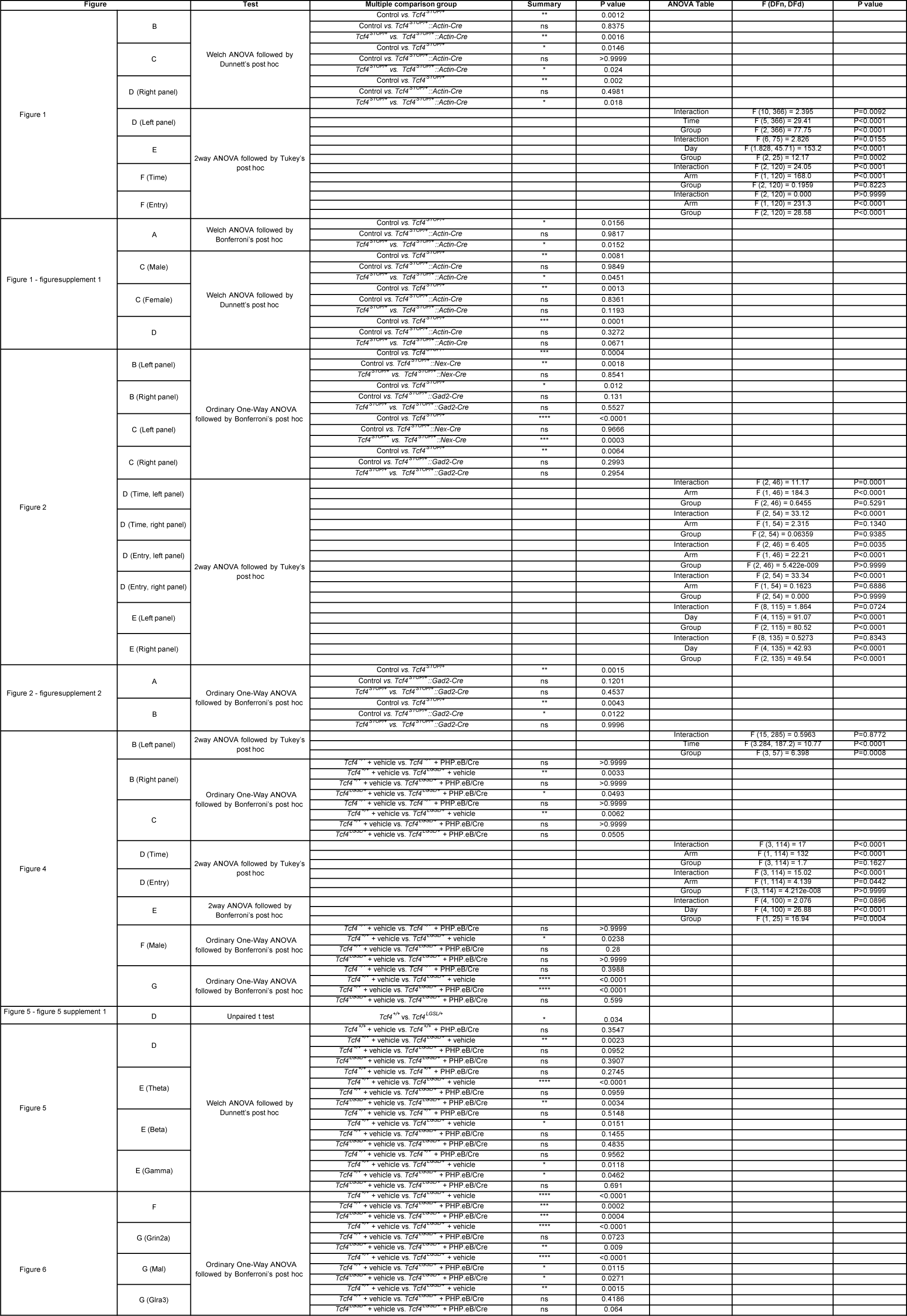
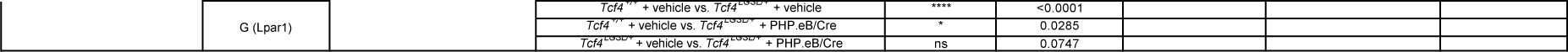
Statistics

## References

1. Amiel, J., Rio, M., de Pontual, L., Redon, R., Malan, V., Boddaert, N., … Colleaux, L. (2007). Mutations in TCF4, encoding a class I basic helix-loop-helix transcription factor, are responsible for Pitt-Hopkins syndrome, a severe epileptic encephalopathy associated with autonomic dysfunction. Am J Hum Genet, 80(5), 988–993. doi:10.1086/515582

2. Bedeschi, M. F., Marangi, G., Calvello, M. R., Ricciardi, S., Leone, F. P. C., Baccarin, M., … Zollino, M. (2017). Impairment of different protein domains causes variable clinical presentation within Pitt-Hopkins syndrome and suggests intragenic molecular syndromology of TCF4. Eur J Med Genet, 60(11), 565–571. doi:10.1016/j.ejmg.2017.08.004

3. Brennand, K. J., Simone, A., Jou, J., Gelboin-Burkhart, C., Tran, N., Sangar, S., … Gage, F. H. (2011). Modelling schizophrenia using human induced pluripotent stem cells. Nature, 473(7346), 221–225. doi:10.1038/nature09915

4. Brzozka, M. M., Radyushkin, K., Wichert, S. P., Ehrenreich, H., & Rossner, M. J. (2010). Cognitive and sensorimotor gating impairments in transgenic mice overexpressing the schizophrenia susceptibility gene Tcf4 in the brain. Biol Psychiatry, 68(1), 33–40. doi:10.1016/j.biopsych.2010.03.015

5. Buzsaki, G., Anastassiou, C. A., & Koch, C. (2012). The origin of extracellular fields and currents--EEG, ECoG, LFP and spikes. Nat Rev Neurosci, 13(6), 407–420. doi:10.1038/nrn3241

6. Deacon, R. M. (2006). Assessing nest building in mice. Nat Protoc, 1(3), 1117–1119. doi:10.1038/nprot.2006.170

7. Dennis, D. J., Han, S., & Schuurmans, C. (2019). bHLH transcription factors in neural development, disease, and reprogramming. Brain Res, 1705, 48–65. doi:10.1016/j.brainres.2018.03.013

8. Deverman, B. E., Ravina, B. M., Bankiewicz, K. S., Paul, S. M., & Sah, D. W. Y. (2018). Gene therapy for neurological disorders: progress and prospects. Nat Rev Drug Discov, 17(10), 767. doi:10.1038/nrd.2018.158

9. Doostparast Torshizi, A., Armoskus, C., Zhang, H., Forrest, M. P., Zhang, S., Souaiaia, T., … Wang, K. (2019). Deconvolution of transcriptional networks identifies TCF4 as a master regulator in schizophrenia. Sci Adv, 5(9), eaau4139. doi:10.1126/sciadv.aau4139

10. Forrest, M., Chapman, R. M., Doyle, A. M., Tinsley, C. L., Waite, A., & Blake, D. J. (2012). Functional analysis of TCF4 missense mutations that cause Pitt-Hopkins syndrome. Hum Mutat, 33(12), 1676–1686. doi:10.1002/humu.22160

11. Forrest, M. P., Waite, A. J., Martin-Rendon, E., & Blake, D. J. (2013). Knockdown of human TCF4 affects multiple signaling pathways involved in cell survival, epithelial to mesenchymal transition and neuronal differentiation. PLoS One, 8(8), e73169. doi:10.1371/journal.pone.0073169

12. Giurgea, I., Missirian, C., Cacciagli, P., Whalen, S., Fredriksen, T., Gaillon, T., … Moncla, A. (2008). TCF4 deletions in Pitt-Hopkins Syndrome. Hum Mutat, 29(11), E242–251. doi:10.1002/humu.20859

13. Goebbels, S., Bormuth, I., Bode, U., Hermanson, O., Schwab, M. H., & Nave, K. A. (2006). Genetic targeting of principal neurons in neocortex and hippocampus of NEX-Cre mice. Genesis, 44(12), 611–621. doi:10.1002/dvg.20256

14. Goodspeed, K., Newsom, C., Morris, M. A., Powell, C., Evans, P., & Golla, S. (2018). Pitt- Hopkins Syndrome: A Review of Current Literature, Clinical Approach, and 23-Patient Case Series. J Child Neurol, 33(3), 233–244. doi:10.1177/0883073817750490

15. Guy, J., Gan, J., Selfridge, J., Cobb, S., & Bird, A. (2007). Reversal of neurological defects in a mouse model of Rett syndrome. Science, 315(5815), 1143–1147. doi:10.1126/science.1138389

16. Hill, M. J., Killick, R., Navarrete, K., Maruszak, A., McLaughlin, G. M., Williams, B. P., & Bray, N. J. (2017). Knockdown of the schizophrenia susceptibility gene TCF4 alters gene expression and proliferation of progenitor cells from the developing human neocortex. J Psychiatry Neurosci, 42(3), 181–188.

17. Jagle, U., Gasser, J. A., Muller, M., & Kinzel, B. (2007). Conditional transgene expression mediated by the mouse beta-actin locus. Genesis, 45(11), 659–666. doi:10.1002/dvg.20342

18. Jung, M., Haberle, B. M., Tschaikowsky, T., Wittmann, M. T., Balta, E. A., Stadler, V. C., … Lie, D. C. (2018). Analysis of the expression pattern of the schizophrenia-risk and intellectual disability gene TCF4 in the developing and adult brain suggests a role in development and plasticity of cortical and hippocampal neurons. Mol Autism, 9, 20. doi:10.1186/s13229-018-0200-1

19. Kennedy, A. J., Rahn, E. J., Paulukaitis, B. S., Savell, K. E., Kordasiewicz, H. B., Wang, J., … Sweatt, J. D. (2016). Tcf4 Regulates Synaptic Plasticity, DNA Methylation, and Memory Function. Cell Rep, 16(10), 2666–2685. doi:10.1016/j.celrep.2016.08.004

20. Kim, H., Berens, N. C., Ochandarena, N. E., & Philpot, B. D. (2020). Region and Cell Type Distribution of TCF4 in the Postnatal Mouse Brain. Front Neuroanat, 14, 42. doi:10.3389/fnana.2020.00042

21. Loo, L., Simon, J. M., Xing, L., McCoy, E. S., Niehaus, J. K., Guo, J., … Zylka, M. J. (2019). Single-cell transcriptomic analysis of mouse neocortical development. Nat Commun, 10(1), 134. doi:10.1038/s41467-018-08079-9

22. Mazur, C., Powers, B., Zasadny, K., Sullivan, J. M., Dimant, H., Kamme, F., … Verma, A. (2019). Brain pharmacology of intrathecal antisense oligonucleotides revealed through multimodal imaging. JCI Insight, 4(20). doi:10.1172/jci.insight.129240

23. Nieuwenhuis, B., Haenzi, B., Hilton, S., Carnicer-Lombarte, A., Hobo, B., Verhaagen, J., & Fawcett, J. W. (2021). Optimization of adeno-associated viral vector-mediated transduction of the corticospinal tract: comparison of four promoters. Gene Ther, 28(1-2), 56–74. doi:10.1038/s41434-020-0169-1

24. Peippo, M. M., Simola, K. O., Valanne, L. K., Larsen, A. T., Kahkonen, M., Auranen, M. P., & Ignatius, J. (2006). Pitt-Hopkins syndrome in two patients and further definition of the phenotype. Clin Dysmorphol, 15(2), 47–54. doi:10.1097/01.mcd.0000184973.14775.32

25. Phan, B. N., Bohlen, J. F., Davis, B. A., Ye, Z., Chen, H. Y., Mayfield, B., … Maher, B. J. (2020). A myelin-related transcriptomic profile is shared by Pitt-Hopkins syndrome models and human autism spectrum disorder. Nat Neurosci. doi:10.1038/s41593-019- 0578-x

26. Quednow, B. B., Brzozka, M. M., & Rossner, M. J. (2014). Transcription factor 4 (TCF4) and schizophrenia: integrating the animal and the human perspective. Cell Mol Life Sci, 71(15), 2815–2835. doi:10.1007/s00018-013-1553-4

27. Rannals, M. D., Hamersky, G. R., Page, S. C., Campbell, M. N., Briley, A., Gallo, R. A., … Maher, B. J. (2016). Psychiatric Risk Gene Transcription Factor 4 Regulates Intrinsic Excitability of Prefrontal Neurons via Repression of SCN10a and KCNQ1. Neuron, 90(1), 43–55. doi:10.1016/j.neuron.2016.02.021

28. Sepp, M., Kannike, K., Eesmaa, A., Urb, M., & Timmusk, T. (2011). Functional diversity of human basic helix-loop-helix transcription factor TCF4 isoforms generated by alternative 5’ exon usage and splicing. PLoS One, 6(7), e22138. doi:10.1371/journal.pone.0022138

29. Sepp, M., Pruunsild, P., & Timmusk, T. (2012). Pitt-Hopkins syndrome-associated mutations in TCF4 lead to variable impairment of the transcription factor function ranging from hypomorphic to dominant-negative effects. Hum Mol Genet, 21(13), 2873–2888. doi:10.1093/hmg/dds112

30. Silva-Santos, S., van Woerden, G. M., Bruinsma, C. F., Mientjes, E., Jolfaei, M. A., Distel, B., … Elgersma, Y. (2015). Ube3a reinstatement identifies distinct developmental windows in a murine Angelman syndrome model. J Clin Invest, 125(5), 2069–2076. doi:10.1172/JCI80554

31. Takano, K., Lyons, M., Moyes, C., Jones, J., & Schwartz, C. E. (2010). Two percent of patients suspected of having Angelman syndrome have TCF4 mutations. Clin Genet, 78(3), 282–288. doi:10.1111/j.1399-0004.2010.01380.x

32. Taniguchi, H., He, M., Wu, P., Kim, S., Paik, R., Sugino, K., … Huang, Z. J. (2011). A resource of Cre driver lines for genetic targeting of GABAergic neurons in cerebral cortex. Neuron, 71(6), 995–1013. doi:10.1016/j.neuron.2011.07.026

33. Thaxton, C., Kloth, A. D., Clark, E. P., Moy, S. S., Chitwood, R. A., & Philpot, B. D. (2018). Common Pathophysiology in Multiple Mouse Models of Pitt-Hopkins Syndrome. J Neurosci, 38(4), 918–936. doi:10.1523/JNEUROSCI.1305-17.2017

34. Wang, F., Flanagan, J., Su, N., Wang, L. C., Bui, S., Nielson, A., … Luo, Y. (2012). RNAscope: a novel in situ RNA analysis platform for formalin-fixed, paraffin-embedded tissues. J Mol Diagn, 14(1), 22–29. doi:10.1016/j.jmoldx.2011.08.002

35. Wang, S., Lai, X., Deng, Y., & Song, Y. (2020). Correlation between mouse age and human age in anti-tumor research: Significance and method establishment. Life Sci, 242, 117242. doi:10.1016/j.lfs.2019.117242

36. Wirgenes, K. V., Sonderby, I. E., Haukvik, U. K., Mattingsdal, M., Tesli, M., Athanasiu, L., … Andreassen, O. A. (2012). TCF4 sequence variants and mRNA levels are associated with neurodevelopmental characteristics in psychotic disorders. Transl Psychiatry, 2, e112. doi:10.1038/tp.2012.39

37. Xia, H., Jahr, F. M., Kim, N. K., Xie, L., Shabalin, A. A., Bryois, J., … McClay, J. L. (2018). Building a schizophrenia genetic network: transcription factor 4 regulates genes involved in neuronal development and schizophrenia risk. Hum Mol Genet, 27(18), 3246–3256. doi:10.1093/hmg/ddy222

38. Xiong, W., Wu, D. M., Xue, Y., Wang, S. K., Chung, M. J., Ji, X., … Cepko, C. L. (2019). AAV cis-regulatory sequences are correlated with ocular toxicity. Proc Natl Acad Sci U S A, 116(12), 5785–5794. doi:10.1073/pnas.1821000116

39. Zeisel, A., Munoz-Manchado, A. B., Codeluppi, S., Lonnerberg, P., La Manno, G., Jureus, A., … Linnarsson, S. (2015). Brain structure. Cell types in the mouse cortex and hippocampus revealed by single-cell RNA-seq. Science, 347(6226), 1138–1142. doi:10.1126/science.aaa1934

40. Zollino, M., Zweier, C., Van Balkom, I. D., Sweetser, D. A., Alaimo, J., Bijlsma, E. K., … Hennekam, R. C. (2019). Diagnosis and management in Pitt-Hopkins syndrome: First international consensus statement. Clin Genet. doi:10.1111/cge.13506

41. Zweier, C., Peippo, M. M., Hoyer, J., Sousa, S., Bottani, A., Clayton-Smith, J., … Rauch, A. (2007). Haploinsufficiency of TCF4 causes syndromal mental retardation with intermittent hyperventilation (Pitt-Hopkins syndrome). Am J Hum Genet, 80(5), 994–1001. doi:10.1086/515583

